# EEG Phase Slips Reveal Detailed Brain Activity Patterns of Novice Vipassana Meditators During Decision Making Tasks

**DOI:** 10.1101/2025.09.03.674031

**Authors:** Ratna Jyothi Kakumanu, Ajay Kumar Nair, Arun Sasidharan, Rahul Venugopal, Ravindra P Nagendra, Bindu Kutty, Ceon Ramon

## Abstract

Our study adopted a novel approach, utilizing four biomarkers, namely EEG potentials, their first-order derivatives, and phase slip rates derived from each, to discern the differences between novice Vipassana (NVP) subjects and non-meditator controls (NMC). Phase slip rates are discontinuities in instantaneous phase that represent cortical phase transitions, indicating a significant change in the overall brain state. We employed 128-channel EEG data from eight NMC and eight NVP subjects, collected during a gamified protocol, to investigate object identification within a visual oddball paradigm. EEG was continuously acquired during the 50 trials for each subject. We retained 44 artifact-free trials per subject (the minimum common across all subjects) and computed within-subject averages. The EEGs were then averaged separately for NMC and NVP subjects. The EEG was filtered in the alpha band, and the phase was extracted using the Hilbert transform, unwrapped, and the phase slip rates were computed. A montage layout of electrodes was used to make the spatiotemporal plots of biomarkers. Our findings revealed that the spatiotemporal profiles of all four biomarkers differed significantly between the two groups of subjects. Furthermore, the NVP subjects demonstrated faster object recognition. These results not only provide a unique method to use phase slip rates to quantify cognitive differences between NMC and NVP subjects but also present a promising set of biomarkers for quantitative task-based EEG analyses.

## A. Introduction

Phase slips derived from the EEG have been extensively studied to understand cortical neurodynamics under various conditions. These conditions include visual object naming tasks, insight moments [1], dynamics of epileptic seizure evolutions [2], and the formation of propagating spatiotemporal phase cone structures on the cortical surfaces [2, 3]. In all of these studies, there is often additional information about the cortical activity not seen in the power spectral density or phase synchronization analysis of EEG data. Our objective in the present study is to investigate whether additional information can be derived from phase slips extracted from the EEG of meditative subjects, particularly for novice Vipassana (NVP) subjects in decision-making tasks. Our results indicate that this approach is feasible, offering a promising avenue for future research in meditation and EEG data analysis.

Phase slips are generally extracted from the EEG data using Hilbert transform techniques. However, we have recently demonstrated that these phase slips can also be extracted from the derivatives of EEG data [4], which provides additional information about cortical neurodynamics compared to the phase slips extracted from EEG alone. Here we have written the first order derivative (d/dt) of EEG as EEG’. The phase slip rates extracted from EEG are written as PSR1, and those extracted from EEG’ as PSR2. In the present study, we examined the spatiotemporal behavior of four quantities, viz., EEG, EEG’, PSR1, and PSR2 of novice Vipassana (NVP) meditation subjects and compared them with the spatiotemporal behavior of non-meditator control (NMC) subjects. Our results indicate there are quantifiable differences between NMC and NVP subjects. In addition, we examined the different stages of visual object recognition between NMC and NVP subjects, as well as the time it takes to recognize an object. A detailed analysis of N1, P1, and P2 visual evoked potentials [5] and their corresponding phase slip rates was performed, and there were noticeable differences between NMC and NVP subjects. The insight moments are also called ‘Eureka’ moments when one recognizes an object or spontaneously finds a solution to a problem [1, 6]. To our knowledge, these results are new and open a new way to quantify the effects of meditation on cortical neurodynamics by use of biomarkers based on phase slip rates, such as PSR1 and PSR2.

In the following sections, we provide a brief overview of the necessary background material on cortical phase transitions, phase slips, Vipassana meditation, and insight moments, which will help clarify our methods and results. Details of the methods, results, and discussions are presented in the subsequent sections.

## B. Background Material

### B.1 Criticality, Phase Transitions, and Phase Slips

Criticality refers to the condition of a system that is finely balanced between order and disorder such that a small perturbation can yield large effects [7]. The sudden large shift in the state of the system is called a phase transition. An example of the critical point is the triple point (273.16 Kelvin, 611.73 Pascal) of water, where it can exist in all three phases of solid, liquid, and gas, and a slight change in the temperature or pressure could cause a shift from one state to the other, e.g., from solid to liquid, etc. Other examples of criticality and phase transitions in nature include superconductivity, ferromagnetism, neuronal activity, and swarm technology [8, 9]. Another typical example of self-organized criticality is the sandpile model [10, 11], where one continually adds sand to a vertical pile until it collapses, triggering cascading avalanches.

Phase slips are discontinuities in rhythmic activity [12, 13] and arise due to the shift from the coordinated synchronous rhythmic activity to more independent asynchronous firing of a group of neurons in the cortex. This generally happens at mesoscopic scales in the brain, but could also occur at larger interconnected areas in the whole brain. The short (i.e., local) and the long (i.e., global) range electrical interconnections in the brain happen over the white matter fiber pathways. Due to these interconnected regions, one can observe phase slips in the invasive ECoG (Electrocorticogram) recordings and noninvasive EEG (Electroencephalogram) recordings. These phase slips represent cortical phase transitions, indicating a significant change in the overall brain state. These changes in brain states could be related to cognitive processes, such as learning or memory, or changes in internal or external stimuli [1, 13]. Examples of internal stimuli include a shift in attention or a change in an internal thought process. Examples of external stimuli include audio or visual stimuli, temperature, pressure, or even social interactions. All of these could cause a phase transition leading toward the formation of a phase slip.

Phase transitions in the brain occur because the electrical activity of the whole cortex is generally in a metastable state, very close to a critical point, and a slight disturbance could cause a phase transition [14, 15]. The phase transitions produce bursts of neural oscillations, which show up as small perturbations in the EEG, MEG, or ECoG data, which becomes a site for the formation of phase slips. Most of these behaviors of criticality in nature can be modeled with an Ising model of ferromagnetism, which has also been applied to the modeling of avalanches in networked neurons [16]. Some other models based on nonlinear many-body field dynamics have also been proposed for the formation of phase slips at mesoscopic scales in the brain [13, 17, 18]. Thus, in summary, one can consider phase slips as a reliable biomarker to study brain behavior, which has been applied to study the cognitive behavior of the brain from EEG [1] and localization of epileptogenic sites from ECoG [3] and EEG [19] data. In the present report, we are applying these phase slip techniques to study the cognitive behavior of novice Vipassana subjects during decision-making processes.

### B.2 Vipassana Meditation and Brain Processes

Vipassana meditation, as outlined in the Pâli Canon, is rooted in the direct experience of the Buddha and the precise method he used to attain enlightenment. As such, it occupies a central place in Buddhist teachings. Vipassana is a comprehensive system of mental cultivation and a transformative process aimed at elevating consciousness [20]. In contemporary neuropsychology, Vipassana is often equated with “mindfulness” and has been widely adopted as a clinical intervention for managing emotional distress, maladaptive behaviors, and psychosomatic conditions [21]. An expanding body of research highlights its efficacy in enhancing psychological and physical well-being [22, 23], making it one of the most extensively studied psychotherapeutic practices globally. The practice has attracted increasing interest in neuroscience for its potential to foster cognitive flexibility, emotional regulation, and psychological resilience [24, 25].

Despite the growing empirical interest, many studies have neglected the underlying philosophical context of meditation and its broader implications for consciousness and well-being [26]. Moreover, existing research often centers on Western adaptations of Vipassana. Studies on incarcerated populations completing the traditional 10-day courses have found reduced recidivism, depression, anxiety, hostility, and substance use [27]. Studies among experienced Vipassana practitioners in the classical tradition of Sayagyi U Ba Khin, as taught by S.N. Goenka, have identified proficiency-specific psycho-neural correlates of well-being [28, 29]. This tradition aims at a radical transformation of consciousness through a sequence of three interdependent meditative techniques designed to accommodate a wide range of temperaments while maintaining internal coherence (Fig. 1).

**Fig. 1.**
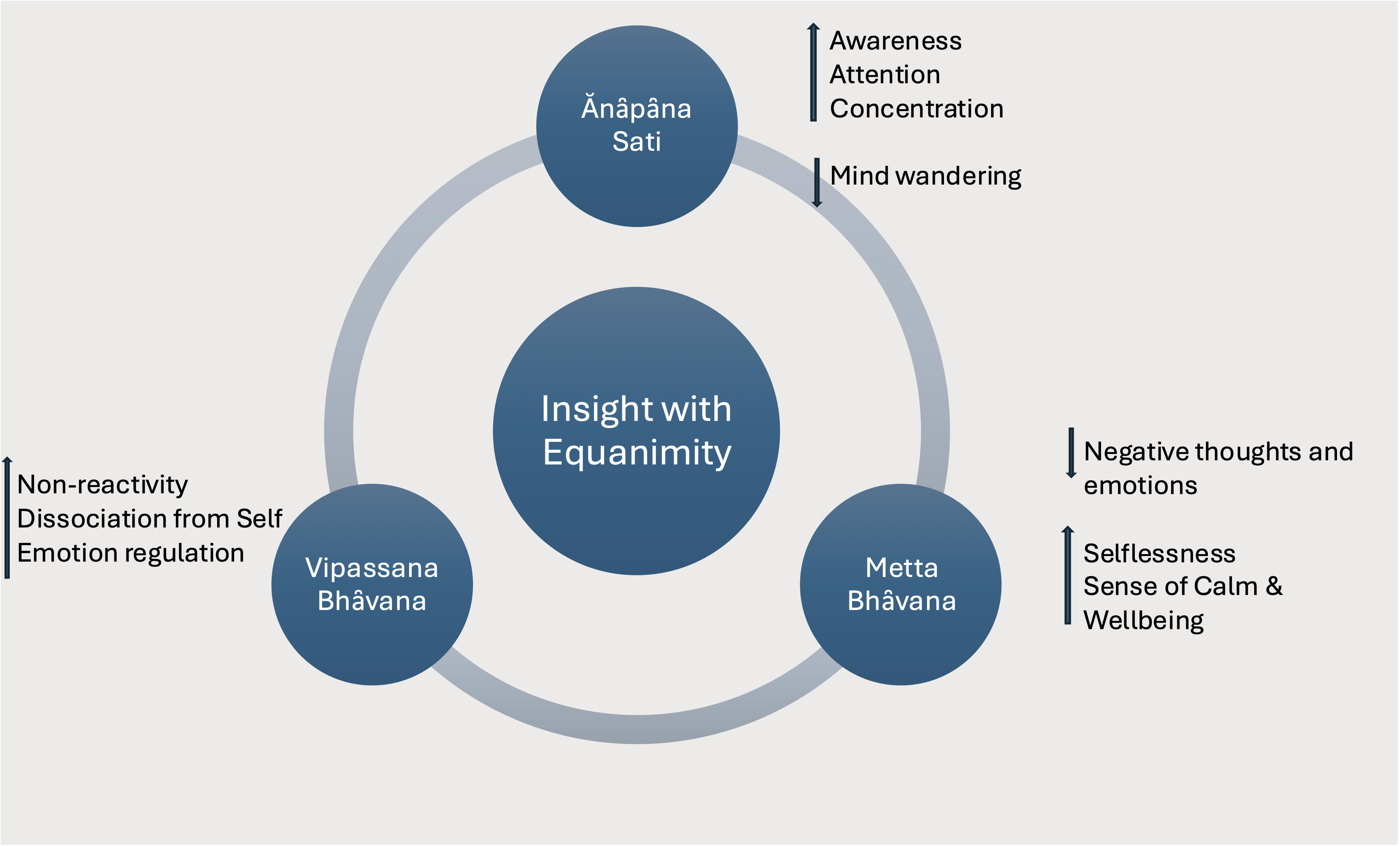
Vipassana Meditation. Schematic representation of the Vipassana practice in the tradition of Sayagyi U Ba Khin as taught by S.N. Goenka.

The training begins with Ânâpâna Sati (Focused Attention Meditation), in which attention is anchored to the breath. This practice cultivates sustained attention, heightened awareness, and perceptual acuity while reducing habitual mind-wandering [30].

Next, during Vipassana Bhâvanâ (Insight Meditation, Mindfulness or Open Monitoring Meditation), practitioners learn to experience the phenomena within the mind-body continuum with objective awareness. These phenomena, including thoughts and feelings, both pleasant and unpleasant, are seen as transient, insubstantial, and ultimately dissatisfactory. Through this sustained, non-reactive observation, the practitioner gains direct insight into the impermanent and impersonal nature of phenomena. Over time, one learns to disidentify from habitual responses of craving and aversion, seen as sources of suffering, and begins to experience increasing inner stability and self-mastery [30, 31, 32].

Finally, Mettâ Bhâvanâ (Loving-Kindness/Compassion Meditation) involves the deliberate cultivation and radiance of goodwill toward oneself and all beings, free from bias or reservation. This phase helps dissolve self-centered patterns and reinforce the equanimity gained through Vipassana, supporting the integration of insight into daily life [30]. Throughout Mettâ practice, the practitioner maintains an objective stance, extending compassionate awareness without expectation.

These three practices are not discrete but are interwoven, each reinforcing the others in a unified framework for inner transformation [33]. Taken together, they engage cognitive, affective, attitudinal, and ethical dimensions, forming a deeply introspective and socially relevant practice [20].

## B.3 Insight Moments and Visual Evoked Potentials

Insight moments are also called ‘Aha!’ or ‘Eureka’ moments, which relate to suddenly or spontaneously finding a solution to a problem. A well-known example is the famous ‘Eureka’ moment of Archimedes. The ‘Eureka’ moment is part of the four stages of creativity suggested by Graham Wallas in 1926, which are commonly called: Preparation, Incubation, Illumination, and Verification [34, 35, 36]. Illumination is the ‘Eureka’ moment. These insight moments can be studied with EEGs, either in the resting state or in evoked response potentials studies, such as looking at a picture on a computer screen while collecting the EEG data [1, 6, 37, 38, 39].

The prominent peaks of visual evoked potentials, P1, N1, and P2, are integral to insight moments, which are generally observed within the first 300 ms from the start of the stimulus onset. After that, within the 0.3 s to 1.2 s time frame from the stimulus onset, several other prominent peaks are also related to different stages of insight moments, allowing the subject to recognize the name, shape, and form of the object on the computer screen and compare them with what is stored in the subject’s memory [1]. Thus, analysis of spatial profiles of EEG and phase slip rates for these peaks and their differences between NMC and NVP subjects is an important topic of investigation. An interesting observation from our results was that there was no remarkable difference in the latency of P1, N1, and P2 peaks for the NMC and NVP subjects. However, there were some noticeable differences in the spatial profiles of these peaks for NMC and NVP subjects. More details are given in the results section.

## C. Materials and Methods

### C.1 Participants

High-density EEG data of 16 healthy adult participants (8 NMC, 8 NVP) were used in this study. No new data sets were collected for this study. We conducted secondary data analyses on a subset of EEG data previously collected in prior projects [28, 29], where details of the data acquisition protocols and participant characteristics have been described. In brief, NMC participants (n=8, 4 females) were in the age range of 23 to 57 years with an average age of 37.6±10 years. For the NVP participants (n=8, 7 females), the age range was 29 to 60 years with an average age of 48.5±10 years. NVP participants had an average meditation experience of 2.2 years with a maximum experience of 3.0 years. The average meditation hours were 1,080. The NMC participants had no prior experience with meditation. These details are also summarized in Table 1.

**Table 1.**
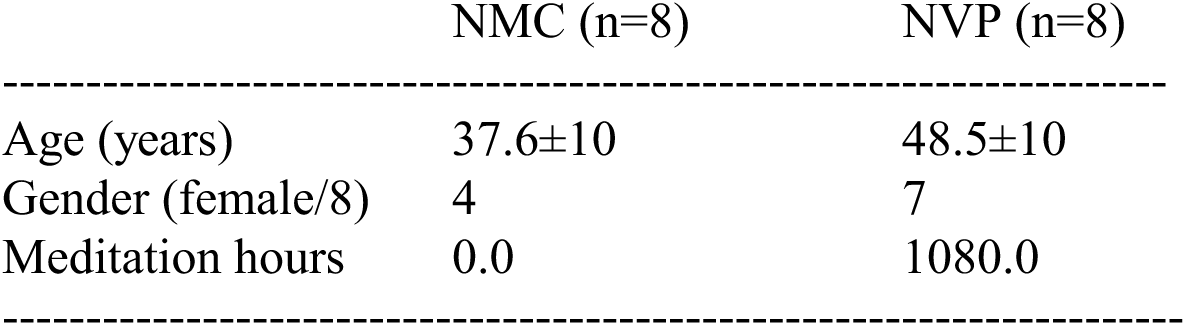
Participant groups’ mean age and estimated hours of meditation experience.

|  | NMC (n=8) | NVP (n=8) |
| --- | --- | --- |
| Age (years) | 37.6±10 | 48.5±10 |
| Gender (female/8) | 4 | 7 |
| Meditation hours | 0.0 | 1080.0 |

The inclusion criteria were: NVP participants should have completed two or three ten-day courses in Vipassana meditation with less than three years of practice and no exposure to any other form of meditation. NMCs should have had no exposure to any form of meditation. All had basic literacy in at least one language: English, Hindi, or Kannada. The exclusion criteria were: participants with neurological or psychological disorders, a history of substance abuse, or on psychiatric or neurological medications. These exclusion criteria were chosen to ensure that the participants’ EEG data were not influenced by any external factors that could impact their brain activity, thereby maintaining the integrity of the study.

Participants were recruited with permission from the Vipassana Research Institute (VRI), Igatpuri, India, via study pamphlets and VRI newsletter advertisements at Vipassana training centers across India. Non-Meditating controls were recruited from the local community in Bangaluru, India. All participants volunteered to participate in the study. Food, accommodation, and travel expenses were offered, along with no other form of financial incentives.

All participants provided written informed consent, which was obtained after a detailed explanation of the study’s purpose, procedures, and potential risks and benefits. The NIMHANS Institute’s Human Ethics Committee approved the consent process, which was in accordance with the Declaration of Helsinki (1964). The study took place at the Human Cognitive Research Laboratory in the Department of Neurophysiology, NIMHANS, Bengaluru, India. Participants had at least a high school education, and the majority belonged to the middle-income category, as defined by Indian standards. Participants were all healthy, right-handed, non-smokers and refrained from any caffeinated beverages on the day of the study.

Although the meditators were recruited from all over India and were practicing a standard form of meditation, no systematic sampling approach, such as cluster sampling, was used to obtain a representative national sample. However, our selected participants represent a typical sample of the educated middle class who may practice Vipassana meditation. As meditation has become a global phenomenon in today’s world, our sample would also represent the trends of modern society in various parts of the world. This global relevance extends to the non-meditating control samples in the middle-class society of India and worldwide.

### C.2 EEG Data of Subjects

A 128-channel EGI system (Electrical Geodesics, Inc., Eugene, OR, USA; now part of Magstim Inc., 1626 Terrace Dr, Roseville, MN 55113) was used for the data collection. Details on the task parameters have been previously reported [40]. In brief, participants engaged in ‘ANGEL’, a gamified audiovisual oddball task with three levels of complexity, presented in 16 blocks each.

The first two blocks of data from the second level (‘Whodunnit’), comprising of 50 trials were used for the present study. Each block had a salient image (black and white face, shape, or a distorted image) presented on one side of a fixation sign, and a non-salient image (black and white checkerboard) presented on the other side. Further, images of one salient image type were presented frequently on one side in a block (for example, a face presented on the right of the fixation) and represented the ‘frequent’ condition. About 20% of the time (‘rare’ trial condition), a salient image of one of the other three image types was presented on the other side (for example, shape to the left of the fixation). Participants were asked to press one of two buttons to indicate if the salient image appeared on the left or the right of the fixation. Figure 2 shows an example of the sequence of images.

**Fig. 2.**
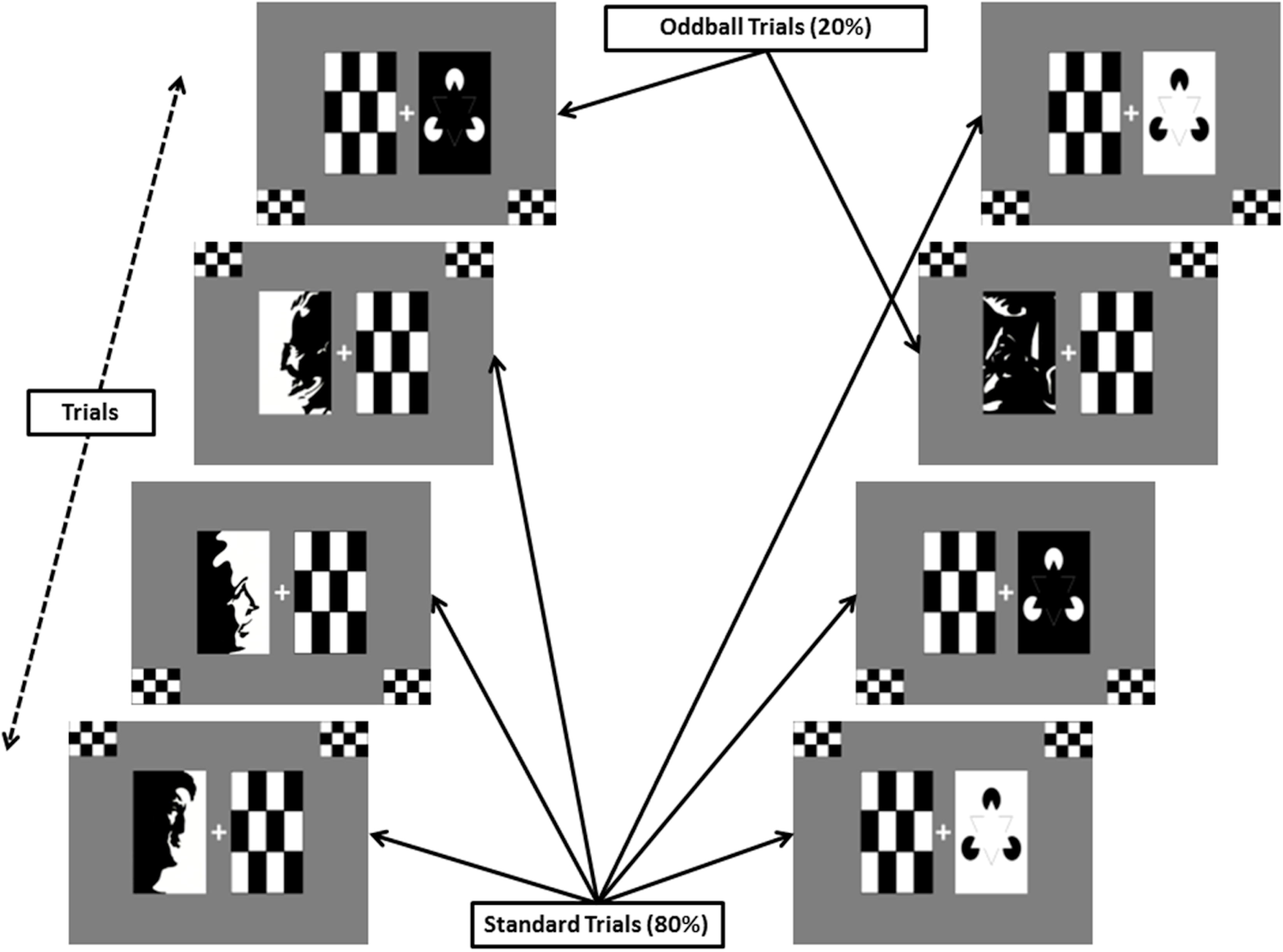
Audiovisual oddball task. Protocol Diagram from earlier publication. Representative examples from two types of task blocks. (From Nair et al., 2016).

### C.3 Preprocessing of the EEG Data

We used MATLAB (MathWorks Inc., Natick, MA, USA) and EEGLAB software 41] for preprocessing the EEG data, phase slip computations, and visualization of results. The data was filtered with an equiripple filter in a 3–49 Hz broadband and then re-referenced to the common averaged reference. The upper frequency of 49 Hz was selected to eliminate the contamination from the power-line frequency of 50 Hz in India, where the data sets were collected. Using the ICA (independent component analysis) techniques, the muscle, eyeblink, and heartbeat artifacts were removed. After that, the data were averaged over trials for each subject. For one subject, we found 44 good trials out of 50. For consistency, we chose 44 good trials out of 50 for all subjects. The averaged trials for eight NMC and eight VPN subjects were averaged. This way, at the end, we had one grand-averaged data set for NMC subjects and another similar data set for NVP subjects. These data sets were then again referenced to the common averaged reference for each channel, filtered in the alpha (7–12 Hz) band with an equiripple filter, and then used for phase slip extraction and PSR computations.

An example of the EEG data after preprocessing is shown in Fig. 3. It is only for two electrodes, one in the frontocentral area (Fig. 3A) and the other in the right visual area (Fig. 3B). Their spatial location is shown in Fig. 3C. The periods of prestimulus, stimulus, and poststimulus period are marked on the top of the EEG plot in Fig. 3A. For analysis and display the time scale for the prestimulus period is −1.0 s to 0.0 s, for the stimulus period is from 0.0 s to 0.240 s, and for the poststimulus period is from 0.240 s to 2.0 s, respectively. The prestimulus period is also commonly referred to as the baseline period. Additionally, we use the terminology ‘poststimulus+’ to refer to the combined duration of the stimulus and poststimulus period from 0.0 s to 2.0 s.

**Fig. 3.**
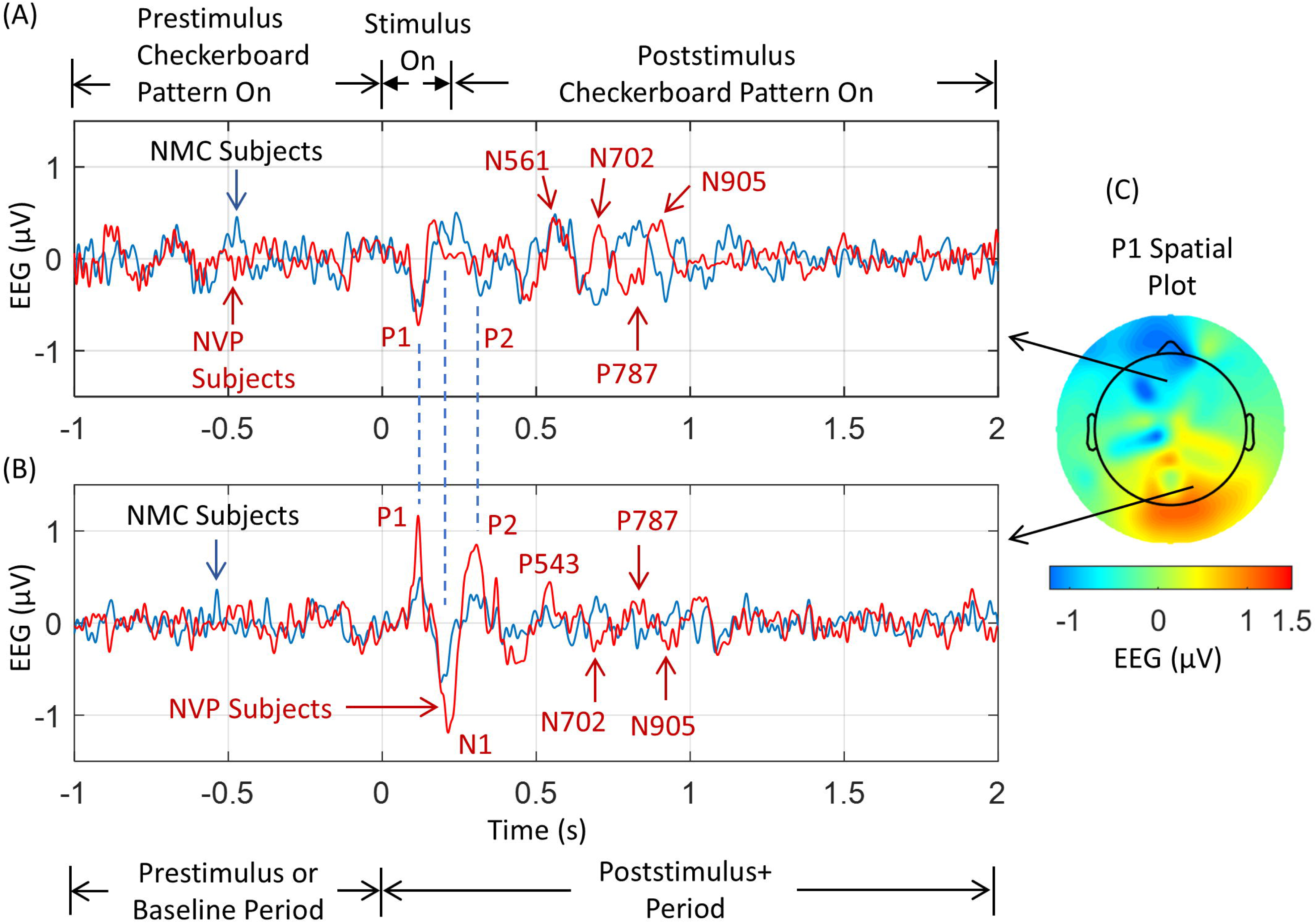
EEG traces and spatial profiles. An example of an EEG trace after preprocessing for two electrodes in a 3-49 Hz broadband. (A) EEG from one of the electrodes in the frontocentral area, (B) EEG from one of the electrodes in the right visual area, and (C) spatial profile of EEG potentials at the P1 peak of the NVP subject. Note that the time series profiles of PSRs at two electrodes are significantly different during the 0.0-1.0 s period, when the subject is visualizing and analyzing the image of the object during covert object naming tasks.

The EEG plot of the right visual electrode (Fig. 3B) has three prominent peaks marked as P1, N1, and P2, which are typical visual event-related potentials [5] observed when one begins to look at an object. It is part of the perception cycle during the stimulus period. The same peaks are also marked in Fig. 3A for the frontocentral electrode. Notice that the polarity of the peaks is reversed in Fig. 3A compared to Fig. 3B. For simplicity, we will use the positive and negative nomenclature of the peaks from the visual electrode in Fig. 3B.

An interesting feature is that these peaks for NMC and NVP subjects are very close. This would suggest two things. First, our data collection and stimulus protocol are robust, so the data collected over several days does not smear the prominent peaks in the EEG data. The second point is that the initial response of the brain to a visual stimulus is very similar in NMC and NVP subjects. For the NVP subjects, these P1, N1, and P2 peaks are located at 115 ms, 214 ms, and 303 ms, respectively, from the start of the stimulus period at 0.0 s. The location of these peaks for the NMC subjects is at 122 ms, 190 ms, and 305 ms, respectively. More details about these peaks are given in the Results section.

During the period of 0.4 to 1.0 s, several prominent peaks with different locations are observed for NMC and NVP subjects. These peaks correspond to the various stages of insight moments [1, 37] related to decision-making and object recognition processes, such as searching for the name and form of the object in memory [1]. These insight moments can be studied during both the stimulus period [6] and/or during the poststimulus period [1, 42, 43]. A detailed analysis of these insight moments for NMC and NVP subjects is given in the Results section.

A note about the spatial plot in Fig. 3C, which will also apply to subsequent spatial plots. The head circle represents the largest possible diameter of the head, as seen from the top, and it covers most of the electrodes on top of the head, positioned slightly above the level of the eyebrows. In addition, there are several lower-level electrodes positioned around the head, which are located below the eyebrows and extending down to ear level. The electrical activity on these electrodes is shown outside the head circle. These electrodes will pick up the electrical activity of the sources in the temporal areas, as well as from the lower, i.e., inferior, and deeper parts of the brain, such as the amygdala, based on the orientation of the sources. There are some electrodes on the forehead, whose electrical activity will be included outside the head circle above the nose in spatial plots. A few of the electrodes on the face are not included in the spatial plots. More details about this can be inferred from the topographic plot script (topoplot.m) included in the EEGLAB software.

### C.4 Phase Slip Rate Computations

The procedures to compute PSR are described in our previous studies [1, 4]. Similar procedures have been used here. The sawtooth patterns of the phase were extracted from the EEG data using the Hilbert transform, which, on unwrapping and detrending, shows phase slips at the locations of small perturbations in the EEG data related to cortical phase transitions. A pictorial representation of phase slip extraction from EEG and its first derivative is given in Fig. 4. It shows the EEG and its first order derivative, EEG’ from one of the electrodes in the front central area, sawtooth pattern of the phase, unwrapped phase with one of the large episodic phase shifts, and several of the large sharp and small broad phase slips. Our interest will be only in the phase slips in the alpha band of 7-12 Hz, and, thus, larger phase slips will not be counted in PSR computations. The PSR1 is from the EEG, and the PSR2 is from the first derivative of the EEG, respectively. Here, EEG voltage is represented as a time series data: *y = y_1_ + y_2_ + + y_n_*, and *y’* is *(dy/dt*) of the time series. The *y’* was computed by the first order differencing of the time series *y* and then dividing it by the digitization interval. A similar representation has been used earlier [4, 44].

**Fig. 4.**
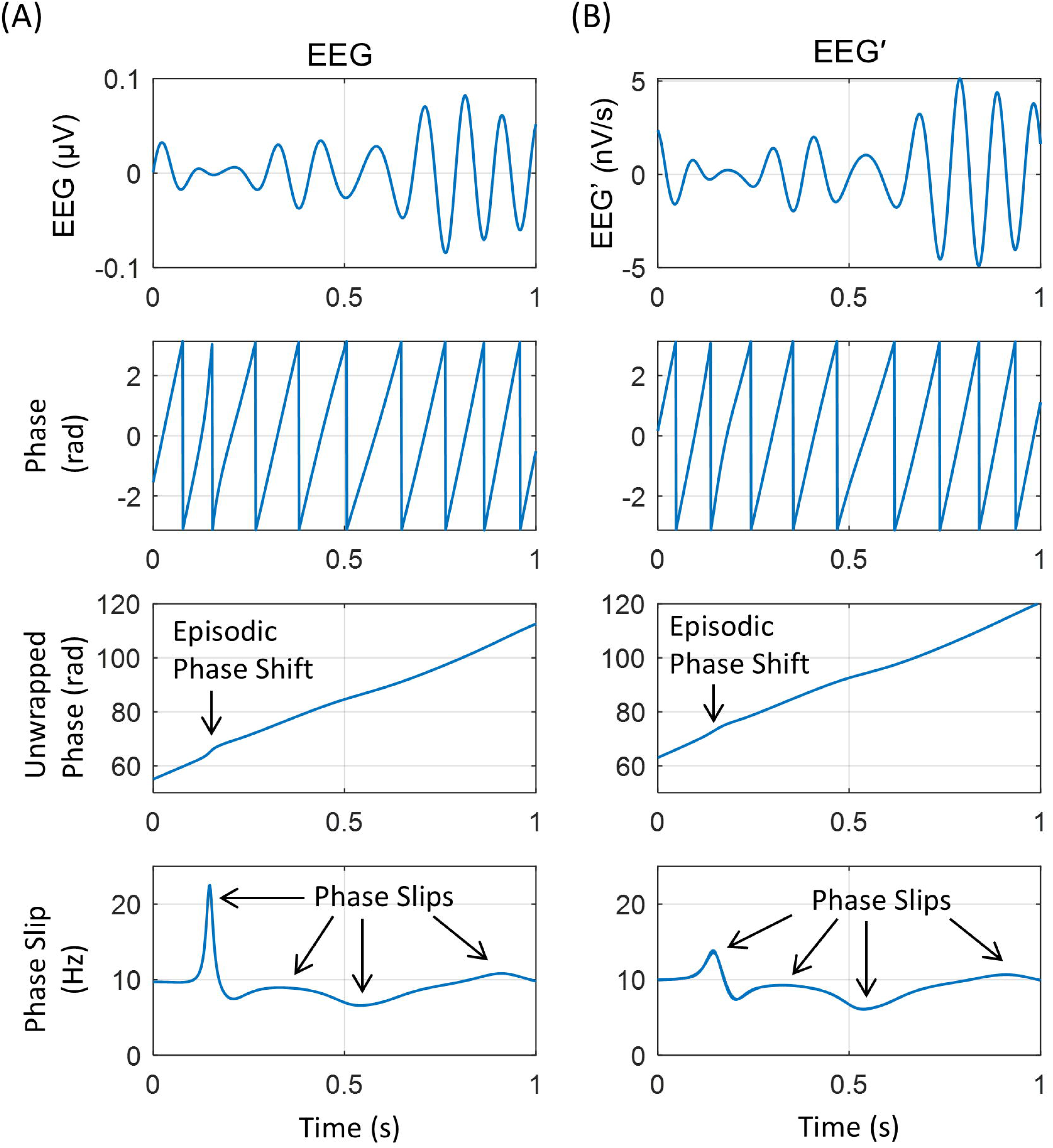
A pictorial representation of the phase slip extraction steps. Plots in the left column are marked in (A) for the phase slips extraction from the EEG, and, similarly, the right column (B) is for the phase slips extraction from EEG’, i.e., from the first order derivative of EEG. The top row is the EEG and EEG’, the second row from the top is the sawtooth pattern of the phase, the third row from the top is the unwrapped phase, and the bottom row shows some of the phase slips.

The PSR from the phase slips was calculated in a stepping window of 10 ms duration with a step size of one digitization point equal to 1.0 ms based on the sampling rate of 1 kHz. The window size is selected based on the Shanon entropy function [45, 46, 47] such that it is short enough to capture the information (lower entropy) but not wide enough to smear (higher entropy) the information. For our case, a window size of 10 ms will give us a maximum value of PSR of 10 counts/10 ms if there were phase slips at each digitization point. In general, it will be difficult to find phase slips at each digitization point, and thus, PSR will be less than 10 counts in each window. The step size of one digitization point, i.e., 1.0 ms, is 10% of the window size, which provides smooth transitions without smearing the information content in going from one stepping window to the next. Also, this window size is approximately 7% of the time period of the lowest frequency (7 Hz) of interest in the alpha band, and the step size is adequate to avoid aliasing errors due to undersampling.

These procedures were used for computing phase slip rates, PSR1 from the EEG data, and PSR2 from the first order derivative of the EEG data. The spatial plots of EEG, PSR1, and PSR2 were constructed with a montage layout of 128 electrode positions on a flat surface.

To separate the phase slips arising from the biological processes versus the random noise, several criteria were applied, which we have used before [4]. These include: (1) phase slip frequency is within the alpha band of 7–12 Hz, (2) sign of the positive or negative peaks did not change for at least three consecutive time steps, and (3) the magnitude of the three consecutive peaks was within the mean ± 1.05ó of the three peaks, where ó is the standard deviation of the mean. The application of these criteria significantly reduced the counting of phase slips due to random noise, while simultaneously maximizing the counting of phase slips from biological processes.

### C.5 Surrogate Data Testing

After filtering in the alpha band, the PSR of random noise was computed from the randomly shuffled EEG data. We used the ‘randperm’ command in the MATLAB software for this. These techniques have been used before by us [1, 2, 4] and also described in a recent paper [48] on surrogate data analysis. The power spectral density of the randomly shuffled EEG data was significantly different from the original EEG data, and it was close to a white noise. The phase slips were extracted from the randomized EEG data and then randomly shuffled. After that, the PSR1 was computed in a stepping window of 10 ms duration with a step size of one digitization point. The mean and standard deviations of PSR1 were found to be zero after averaging over n = 100 trials. A similar analysis was also performed on the PSR2 derived from the first order derivatives of the randomly shuffled EEG data. It was found that in this case also, the mean and standard deviations of PSR2 from the shuffled data were zero after averaging over n = 100 trials. This shows that our reported results are from the biological processes extracted from the EEG data and are above the results from the random noise.

### C.6 Statistical Analysis

To compare two data sets, we used a paired Student’s t-test or histogram distributions. However, these two tests often fail if the means and standard deviations of two distributions are very close to each other. Similarly, the Kolmogorov-Smirnov test also fails if two cumulative distribution functions are similar, even though there are observable and quantifiable differences in their spatial plots. In such situations, the Relative Difference Measure (*RDM\**) and the Magnification Factor (*MAG*) are considered to be better choices than the Student’s t-test, the Kolmogorov-Smirnov test, or the histogram distribution to compare two spatial plots of similar values [2, 45]. It is a prevalent technique to quantify the differences between two spatial plots of data, such as in EEG spatial plots [2, 45]. Most of our results are spatial plots of EEG or phase slip rates. The *RDM\** is defined as:

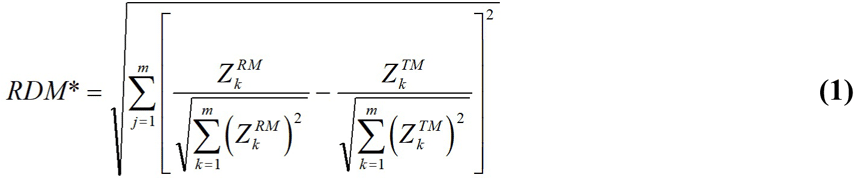

where index, *j=1:m*, runs over all 128 electrodes. The *Z_k_ ^RM^* are the values for the reference measurement, and *Z_k_ ^TM^* are the values of the test measurements. As an example, it could be the EEG potential at the *k^th^* electrode for the reference measurements (RM) and the test measurements (TM), respectively. The magnification factor is defined as:

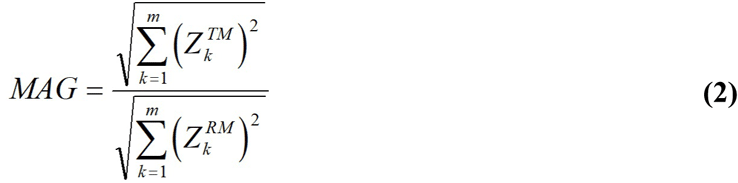

The *RDM\** is a measure of the difference in spatial profiles of two data sets, and *MAG* is a measure of differences in the magnitude of the two data sets. If two data sets are the same, *RDM\** will be zero and *MAG* will be unity. The degree of differences is quantified with higher values of *RDM\** in the range of 0.0 to 1.0. The maximum value of *RDM\** is 1.0 if the test measurements are zero. The variation in the value of *MAG* from unity will quantify the differences in the magnitude of the two data sets. Values of *MAG* greater than 1.0 would mean that the test measurements (TM) are higher in magnitude as compared with the reference measurements (RM). And the reverse is true if *MAG* is less than 1.0.

To compare the means of more than two data sets, we used ANOVA (Analysis of Variance). Histogram plots of data sets were also made. Mean and standard deviations of two histogram plots were computed and compared. In addition, two histogram distributions were also compared with their RMS bin count differences, *RMS H_diff_*, defined as:

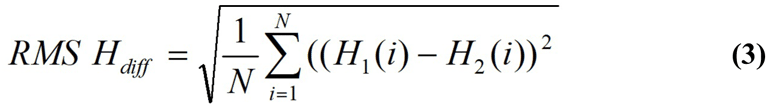

where *H_1_*and *H_2_* are two histogram distributions with *N* bins.

### C.7 Computational Resources

The data analysis was performed on a desktop computer with an 8-core CPU and 112 GB of memory using EEGLAB [41]. For computations of PSR requiring more significant (>112 GB) memory, we used MATLAB software at the UCSD (University of California, San Diego) supercomputing center using their NSG (Neuroscience Gateway) portal.

## D. Results

### D.1 Non-meditator Control (NMC) Subjects

The temporal plots of EEG, EEG’, PSR1, and PSR2 of one of the electrodes in the front central area are given in Fig. 5. During the stimulus and poststimulus periods, the prominent event-related potential peaks are recognizable in the top left plot of broadband EEG in the 3-49 Hz, which are difficult to see in the alpha band EEG plot, (left and right middle plots). The prominent peaks in the EEG are difficult to recognize in the plot of the EEG’ (top right plot).

**Fig. 5.**
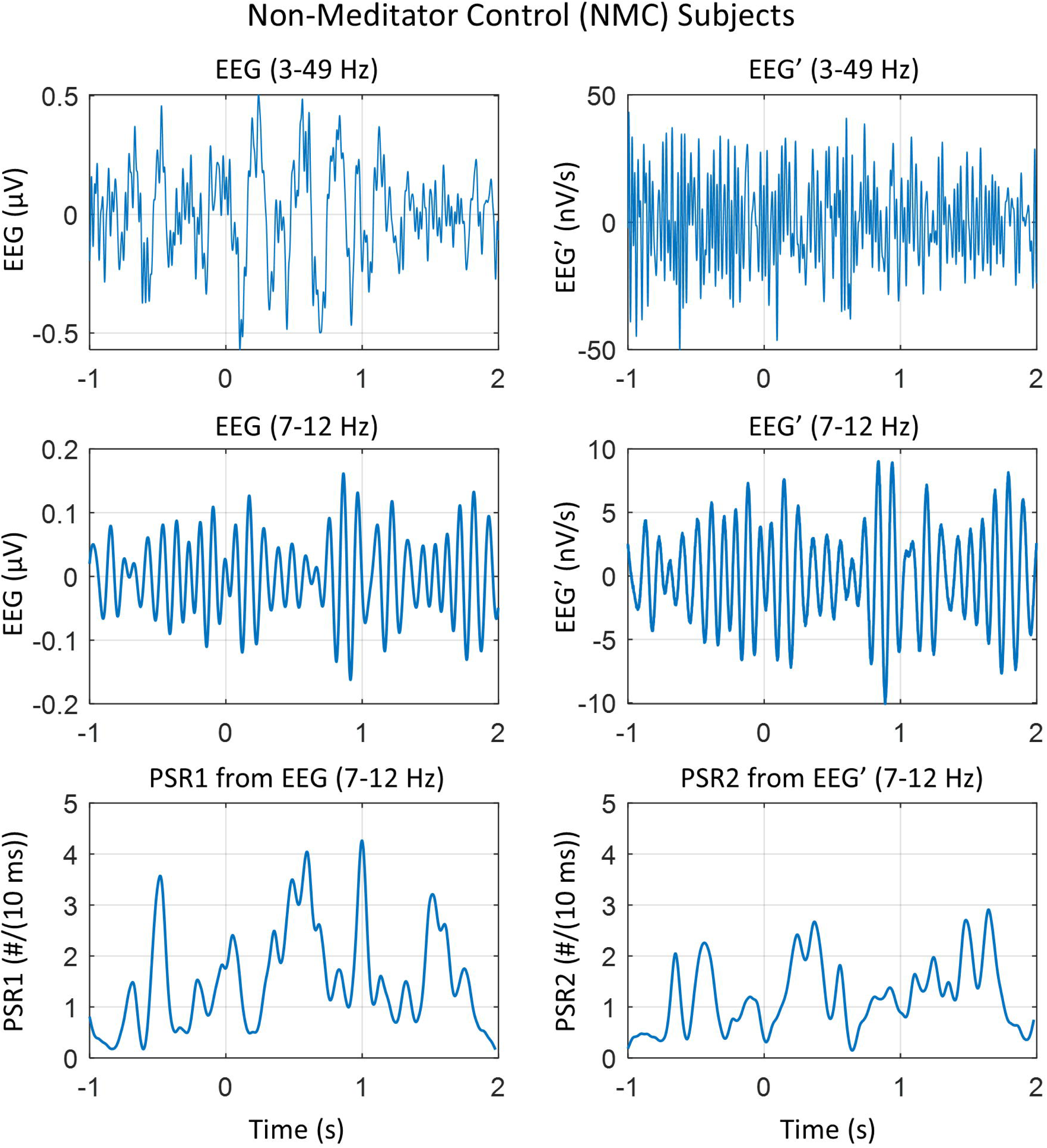
EEG, its derivative, and phase slip rates. Temporal plots of quantities of one of the electrodes in the front central area for non-meditator control (NMC) subjects. (Top left) EEG in 3-49 Hz band, (top right) EEG’ in 3-49 Hz band, (middle left) EEG filtered in the alpha band of 7-12 Hz, (middle right) EEG’ filtered in the alpha band of 7-12 Hz, (bottom left) phase slip rate PSR1 extracted from EEG in the alpha band, and (bottom right) phase slip rate PSR2 extracted from EEG’ in the alpha band. Notice the prominent peaks in PSR1 and PSR2 related to brain activity during the visual object recognition tasks. The prestimulus period is from −1.0 to 0.0 s. The combined stimulus and poststimulus period, referred to as the poststimulus+ period, spans from 0.0 to 2.0 seconds.

They tend to disappear with the first-order differentiation of the EEG. However, the small high-frequency perturbations in the EEG are amplified due to the removal of linear trends through differentiation. In the alpha band, one can see the presence of the alpha wave spindling and small perturbations in the EEG and also in the EEG’.

The PSR1, computed from the EEG data filtered in the alpha (7-12 Hz) band, provides crucial insights. Notably, the PSR1 plot reveals a small and large peak between −0.9 s and −0.45 s during the prestimulus period. These peaks are associated with the background activity of the brain while looking at the checkerboard pattern during the prestimulus period. The broad peak during the poststimulus period, stretching from 0.15 s to 0.95 s, with the prominent peak centered at 0.6 s, is equally significant. This peak is related to the initial response of the visual cortex and the recognition of the object in the language and memory areas of the brain, a common occurrence in the insight moment analysis of visual object naming [1]. The broad peak between 1.35 and 1.9 s is probably related to the brain returning to the background activity after the end of visual object naming processes.

The plot of PSR2 reveals the rate of change in the neuronal population participating in the formation of scalp EEG. It exhibits a bifurcated peak between −0.8 and −0.2 s during the prestimulus period, which correlates with the background activity of the brain while viewing the checkerboard pattern. During the first second of the combined stimulus and poststimulus period, a broad peak is observed from 0.0 to 0.8 s, with the main peak at 0.45 s, which is likely related to the recognition of the object in memory and language areas [1, 37]. The several peaks from 1.0 s to 2.0 s correspond to the return to background activity.

The spatial plots of the root mean square (RMS) of EEG and EEG’, and mean values of PSR1 and PSR2 for NMC subjects are given in Fig. 6. These values were computed within a stepping window of 0.5 s covering the period of −1.0 to 2.0 s.

**Fig. 6.**
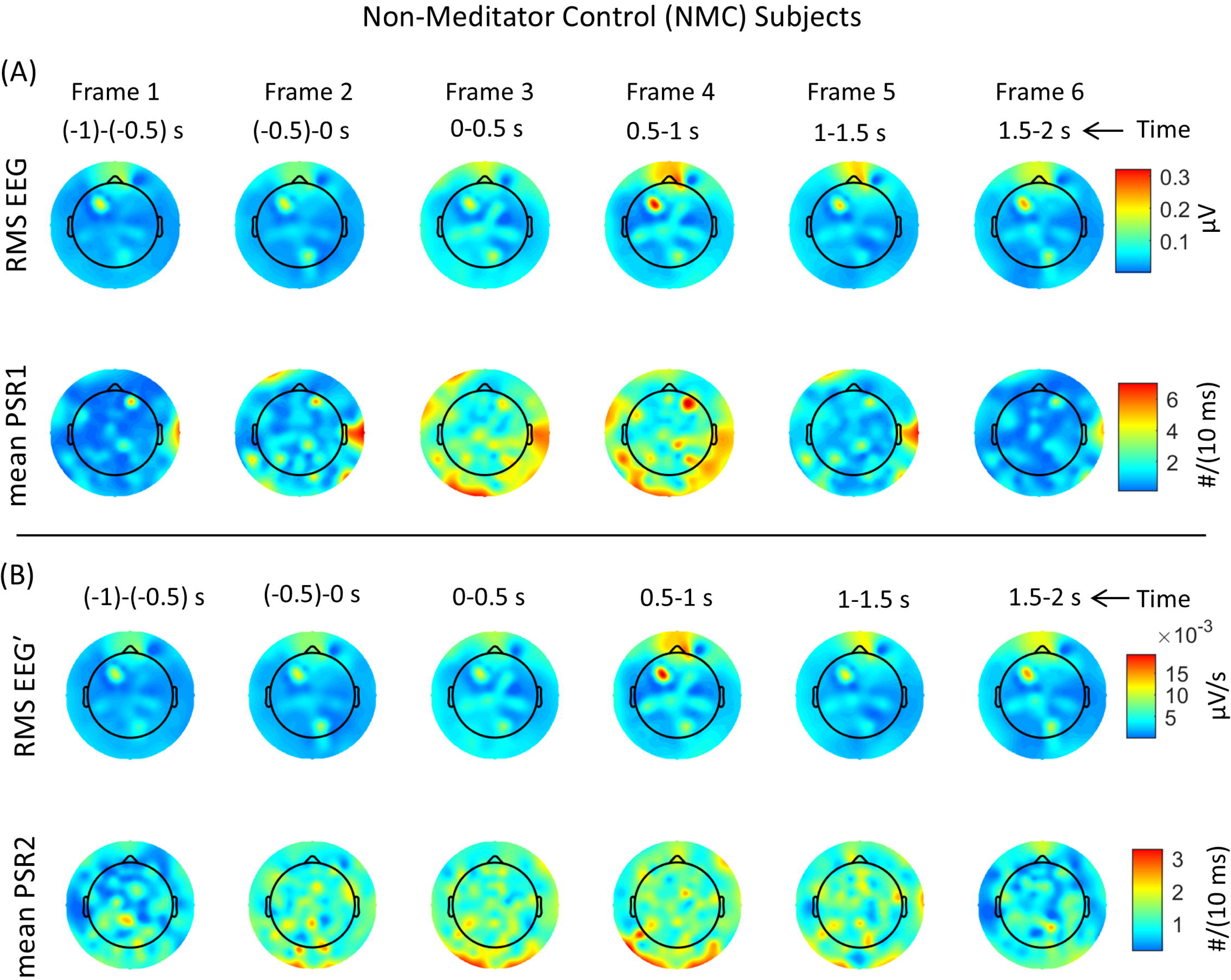
Plots of EEG and PSR of non-meditator control (NMC) subjects. Spatiotemporal plots of the root mean square (RMS) values of EEG and its first order derivative (EEG’) and their associated mean values of phase slip rates for non-meditator control (NMC) subjects. The RMS and mean values were computed within a 0.5 s stepping window. The temporal duration for each frame is given at the top. (A) The top two rows are for the RMS EEG and the mean PSR1, respectively. (B) The bottom two rows are for the RMS EEG’ and the mean PSR2. Notice the shifting spatial patterns of all four quantities.

Fig. 6 is divided into sections (A) and (B). The top two rows in section (A) present the spatial plots of RMS EEG and the mean values of associated phase slip rate, i.e., mean of PSR1 (#/(10 ms)), respectively. The RMS and mean values were computed within a 0.5-s stepping window. Similarly, the bottom two rows marked as section (B) are the spatial plots of the RMS value of the first-order derivative of EEG, i.e., EEG’, and the mean values of its associated phase slip rate, PSR2 (#/(10 ms)), respectively. The frame numbers and their temporal durations are given at the top. The first two frames, i.e., Frame 1 and Frame 2, are during the prestimulus period. The Frame 3 from 0.0 s to 0.5 s is for the combined period of stimulus (0.0 to 0.240 s) and part of the poststimulus (0.240 s to 0.5 s) period. The remaining three frames are for the poststimulus period.

In all frames, RMS EEG activity is intense in the forehead area above the nasion point, corresponding to the brain activity in the brain’s frontal area responsible for the emotional responses to external stimuli, such as visual images on the computer screen. In Frame 1 and Frame 2, it is subdued, begins to increase in Frame 3, and becomes strongest in Frame 4. After that, it starts to decrease in subsequent Frames 5 and 6. There is another activity spot in the left frontocentral area, which is present in all six frames. It is of moderate strength (0.2 µV) during the prestimulus period but becomes stronger during the poststimulus period, with the most substantial value of 0.3 µV in Frame 4. The mean PSR1 plots (Fig. 6A, bottom row) differ from RMS EEG plots. There is a bright spot near the right ear in the central right temporal area. It is present during the stimulus period and also during the poststimulus period. Its activity is noticeable in Frames 3 to 4 during the poststimulus period and almost disappears in Frame 6.

There is also an active spot in the right frontocentral area, located just to the right side, below the nose. The activity at this spot is high in Frames 3 and 4, which matches the several peaks of event-related potentials in the EEG during this period (Fig. 5, top left plot). An interesting feature to note is that mean PSR1 activity is widespread in the lower half of the plot, with some areas in the central and parietal regions, as well as others near the left and right edges of the plot. These corresponding features are not visible in the spatial plots of the RMS EEG.

The spatial plots of RMS EEG’ and the corresponding mean PSR2 are given in Fig. 6B. The spatial features in RMS EEG and RMS EEG’ are very similar, which is expected because EEG’ is the first order derivative of the EEG. Similarly, the spatial patterns of mean PSR2 are similar to those of mean PSR1, representing the rate of change of the neuronal population involved in cortical electrical activity.

A one-way ANOVA analysis was performed on the six frames of RMS EEG, and it was found that the mean of all six frames were significantly (p < 0.01) different from each other. A similar analysis was performed on the remaining three quantities, namely, RMS EEG’, mean PSR1, and mean PSR2. Results were similar, i.e., the means of the six frames for each quantity were significantly (p < 0.01) different from one another. A comparative analysis of the six spatial plots of EEG and PSR1 was done with the paired Student’s t-test, and it was found that each frame was significantly (p < 0.01) different from the others. A similar analysis was performed on each frame of RMS EEG’ and the mean PSR2, and it was found that they were significantly (p < 0.01) different from each other.

### D.3 Novice Vipassana (NVP) Subjects

All quantities related to one of the frontocentral electrodes for NVP subjects are given in Fig. 7. There are well-defined and recognizable peaks in the EEG during the prestimulus, stimulus, and poststimulus periods. During the prestimulus period, the prominent peaks are located at −0.88 s, −0.793 s, and −0.67 s. These peaks are related to the background activity of looking at the checkerboard pattern on the computer screen. During the stimulus and poststimulus periods, the prominent peaks are at: 0.172 s, 0.4 s, 0.56 s, 0.702 s, and 0.903 s. These peaks are related to insight moments [37] for visualization and recognition of the object during the stimulus and poststimulus periods [1]. Their temporal location for NVP subjects differs from that of the NMC subjects. This is also reflected in the plot of PSR1. The prominent peaks are at −0.544 s, 0.436 s, 0.65 s, 0.823 s, and 1.503 s. These plots suggest that the timings for the object recognition steps for NVP subjects are slightly different from those of NMC subjects.

**Fig. 7.**
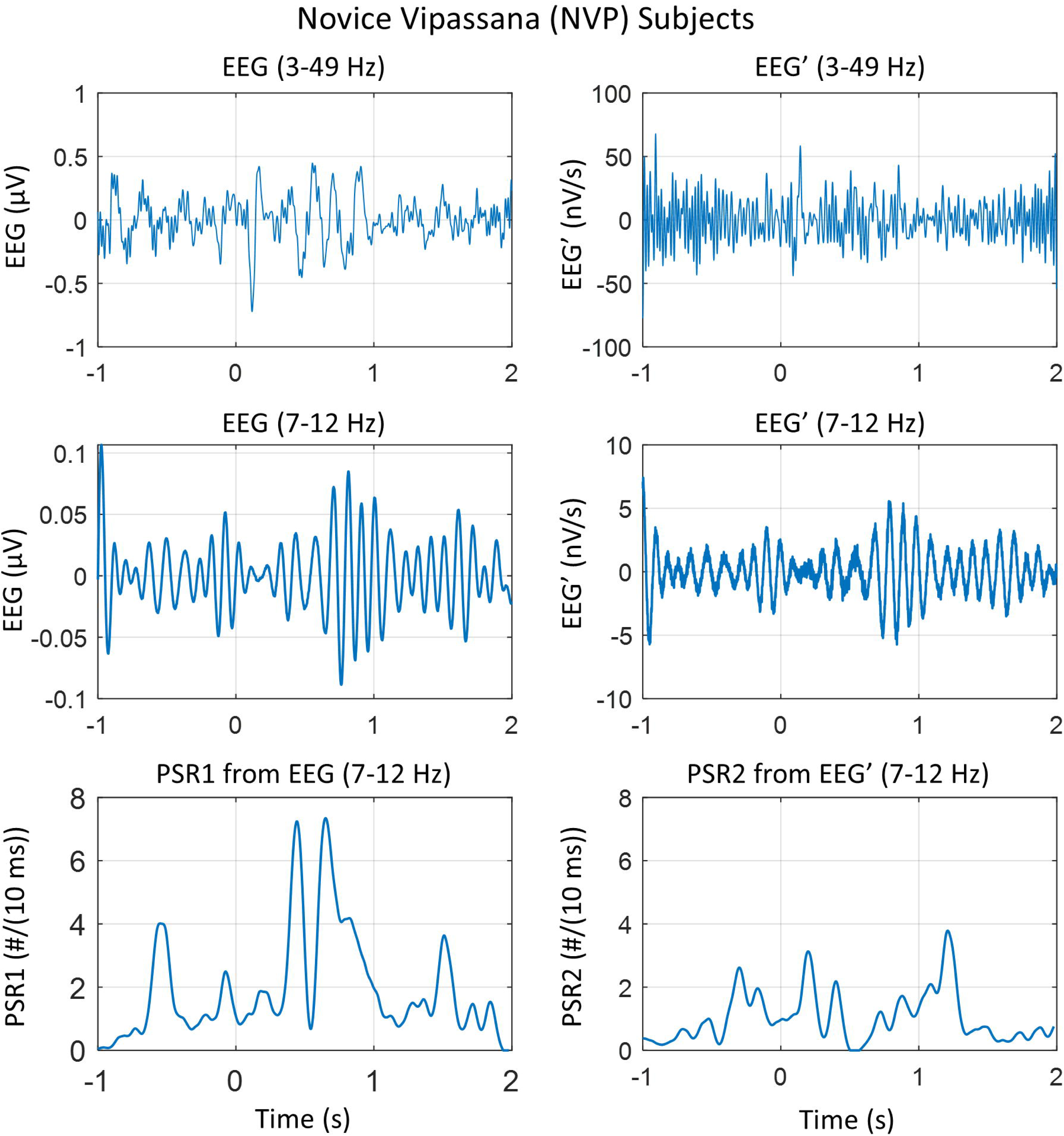
One electrode plot of NVP subjects. Temporal plots of quantities of one of the electrodes in the front central area for novice Vipassana (NVP) subjects. (Top left) EEG in 3-49 Hz band, (top right) EEG’ in 3-49 Hz band, (middle left) EEG filtered in the alpha band of 7-12 Hz, (middle right) EEG’ filtered in the alpha band of 7-12 Hz, (bottom left) phase slip rate PSR1 extracted from EEG in the alpha band, and (bottom right) phase slip rate PSR2 extracted from EEG’ in the alpha band. Notice the prominent peaks in PSR1 and PSR2 related to brain activity during the visual object recognition tasks.

The spatiotemporal plots of brain activity for NVP subjects are given in Fig. 8. These plots are slightly different from the corresponding plots of NMC subjects. There is a bright spot near the nose in the RMS EEG plot in Frame 1. The activity related to the orbital brain is picked up by the electrodes on the forehead above the nasion point. This disappears in the subsequent frames. In Frame 3, there is a large bright spot at the bottom, which relates to the first impressions of the visual stimulus during the stimulus and the start of the poststimulus period. In the following two frames, frames 4 and 5, a low level (∼ 0.1 µV) RMS EEG activity is spread in the broader area in the left frontal, central, and left central parietal areas. These are probably related to a search for the name and form of the stimulus object in the language and memory areas of the brain. Frame 6 represents the return to the brain’s background activity while looking at the checkerboard pattern.

**Fig. 8.**
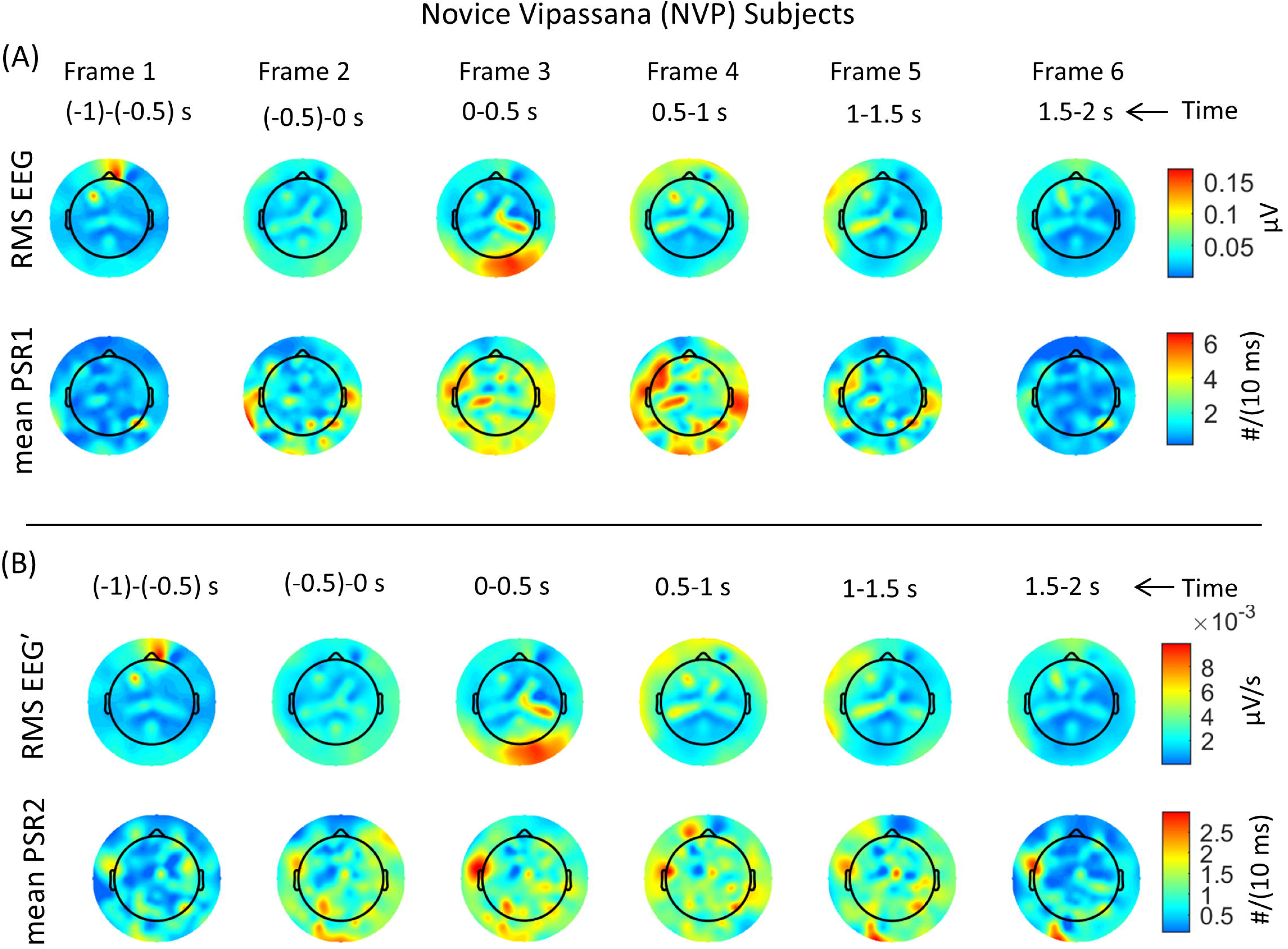
Plots of EEG, its derivative, and PSR of NVP subjects. Spatiotemporal plots of the root mean square (RMS) values of EEG and EEG’ and their associated mean values of phase slip rates for novice Vipassana (NVP) subjects. The RMS and mean values were computed in the stepping window of 0.5 s duration. The temporal duration for each frame is given at the top. (A) The top two rows are for the RMS EEG and the mean PSR1, and (B) the bottom two rows are for the RMS EEG’ and the mean PSR2, respectively. Notice the shifting spatial patterns of all four quantities from Frame 1 to Frame 6.

The spatial patterns of mean PSR1 plots are different from the RMS EEG spatial plots. This is particularly noticeable in frames 3, 4, and 5 when the bulk of the brain activity happens to recognize the visual stimuli on the screen. The brain activity is widespread in different parts of the brain and becomes less in Frame 5. In Frame 6, the PSR1 activity is similar to that in Frame 1, signifying that brain activity has returned to the background level at the end of the trial. The spatial patterns of RMS EEG and RMS EEG’ are similar. Similarly, the spatial patterns of mean PSR1 and PSR2 are very similar except in Frame 6. The PSR2 activity has two bright spots, one near the left central temporal area and one in the left posterior area, which are absent in the PSR1 spatial plot.

A one-way ANOVA analysis was performed on the six frames of four quantities, viz., RMS EEG, RMS EEG’, PSR1, and PSR2. It was found that the means of all six frames were significantly (p < 0.01) different from each other for each of the four quantities. A comparative analysis of the six spatial plots of EEG and PSR1 shown in Fig. 8 for NVP subjects was performed using the paired Student’s t-test, and it was found that each frame was significantly different from the others (p < 0.01). A similar analysis was performed on each frame of RMS EEG’ and the mean PSR2, and it was found that they were also significantly different from each other (p < 0.01).

### D.3 Differences between NMC and NVP Subjects

#### D.3.1 EEG and PSR1 Differences between NMC and NVP Subjects

The poststimulus+ values above the prestimulus, i.e., baseline values, were computed for the NMC and NVP subjects. For these purposes, the stimulus period of 0 to 240 ms was combined with the poststimulus from 0.24 s to 2.0 s, and it is called the poststimulus+ period. In general, the majority of the brain response happens between 0.04 s and 1.2 s from the start of the visual stimulus at 0.0 s [1]. The RMS EEG was computed for the pre- and poststimulus+ periods separately. Then the prestimulus values were subtracted from the poststimulus+ values to obtain the values above the baseline, i.e., above the prestimulus values. The same procedures were repeated for RMS EEG’, mean PSR1, and PSR2.

RMS EEG and mean PSR1 above the prestimulus are given in Fig. 9. The spatial plots of RMS EEG are shown in Fig. 9A, and the mean PSR1 are given in Fig. 9B. The left plot is for NMC subjects, and the right plot is for NVP subjects. For NMC subjects, the RMS EEG activity is very strong at the top of the plots, which is from the electrodes on the forehead area and near the eyes. We have removed the artifacts related to eye motion and eye blinks; thus, these electrodes likely show the activity from the orbital and frontal lobes of the brain. In comparison, the frontal activity is low for NVP subjects. In addition, for NVP subjects, there is more activity outside the head circle in the left temporal areas. Inside the head circle for NMC subjects, there are bright spots in the frontal and left central regions, which are less intense for NVP subjects. The paired Student’s t-test showed a significant (p < 0.01) difference between the spatial plots of the RMS EEG of NMC and NVP subjects. Similarly, the RDM* and MAG values of 1.36 and 1.04, respectively, suggest some noticeable differences between the two spatial plots. The RDM* of 1.36 would suggest a 36% difference in spatial distribution. The MAG of 1.04 would indicate that there is a 4% difference in the magnitude between the two plots. This is very much evident in the differences in the relative magnitude of bright spots at similar locations in the two plots. These differences are also visible in the histogram plot of Fig. 9C. The mean values of histogram counts are the same for both groups, but the standard deviations, ó, are different. The RMS values of histogram bin counts for NMC and NVP subjects are 22.33 and 19.49 counts, respectively. The RMS values of histogram bin counts for NMC and NVP subjects are 22.33 and 19.49 counts, respectively. The RMS difference of histogram bin counts, RMS *H_diff_* is 17.55 counts. This translates to about a 42% difference for the total bin counts, and could be considered a significant difference between the NMC and NVP subjects.

**Fig. 9.**
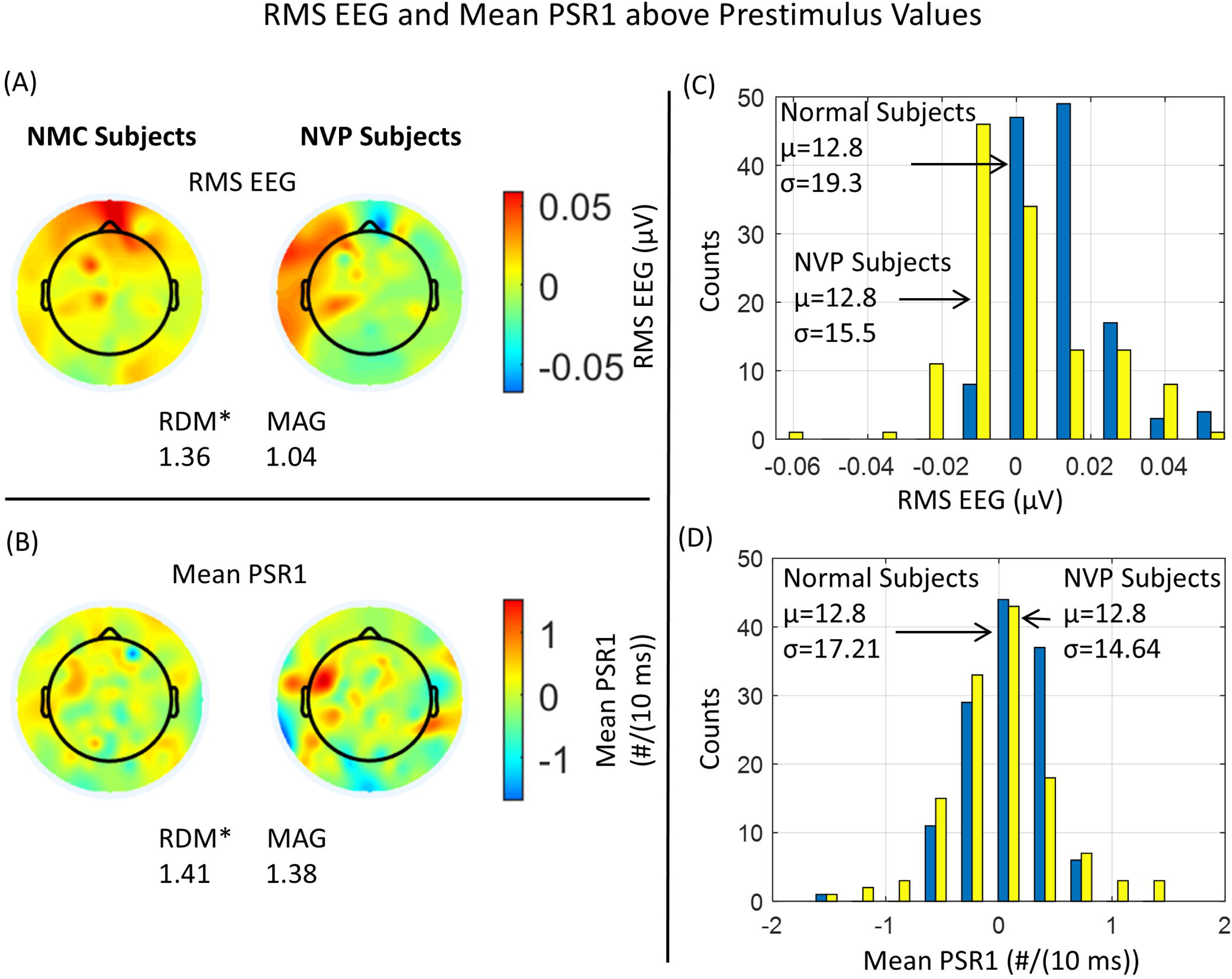
Activity patterns of EEG and PSR1 above baseline. (A) Spatial plots of RMS EEG above the baseline of NMC and NVP subjects, (B) corresponding spatial plots of mean PSR1 above the baseline, (C) histogram distribution of RMS EEG values shown in spatial plots of A, and (D) histogram distribution of PSR1values shown in spatial plots of B.

The spatial plots of PSR1 are given in Fig. 9B, and the corresponding histogram distributions are given in Fig. 9D. For NMC subjects, the mean PSR1 is distributed all over the plot. In contrast, for the NVP subjects, it is strong in the left and right temporal areas. Also, a bright spot near the right ear, which does not have a corresponding high activity in the EEG plot. A paired Student’s t-test was performed on the absolute values of mean PSR1 for the NMC and NVP subjects, and it was found that they were significantly (p < 0.05) different from each other. The *RDM\** and *MAG* values are 1.41 and 1.38, which will also suggest that the two plots are significantly different.

The mean values of histogram counts of PSR1 are the same for both groups, but the standard deviations, ó, are different. The RMS values of histogram bin counts of PSR1 for NMC and NVP subjects are 20.74 and 18.88 counts, respectively. The RMS difference of histogram bin counts is 6.52 counts. This translates to about a 16% difference when normalized to the total number of bins, and could be considered a noticeable difference in the histogram counts of PSR1 between the NMC and NVP subjects.

#### D.3.2 EEG**’** and PSR2 Differences between NMC and NVP Subjects

RMS EEG’, mean PSR2 above the prestimulus values, and their corresponding histogram distributions are given in Fig. 10. For NMC subjects, the RMS EEG’ activity is very strong at the top of the plots, which is from the electrodes on the forehead area and near the eyes. In contrast, for NVP subjects, the RMS EEG’ activity is in the left side temporal regions of the plots. The differences in their spatial distributions are exemplified by the *RDM\** value of 1.27, while their magnitudes are very similar, as indicated by the *MAG* value of 1.08. A paired Student’s t-test was performed on the RMS EEG’ values for the NMC and NVP subjects, and it was found that they were significantly (p < 0.01) different from each other. A histogram distribution of these values are given in Fig. 10C. The mean values of histogram counts are the same for both groups, but the standard deviations, ó, are different. The ó is ±18.56 and ±13.65 for the NMC and NVP subjects, respectively. This translates to about a 26% difference between ó for the NMC and NVP subjects and could be considered a significant difference between the two ó values. The RMS values of histogram bin counts for NMC and NVP subjects are 21.79 counts and 18.21 counts, respectively. The RMS difference of histogram bin counts, RMS *H_diff_* is 16.95 counts. This translates to about a 42% difference for the total bin counts, and could be considered a significant difference between the NMC and NVP subjects.

**Fig. 10.**
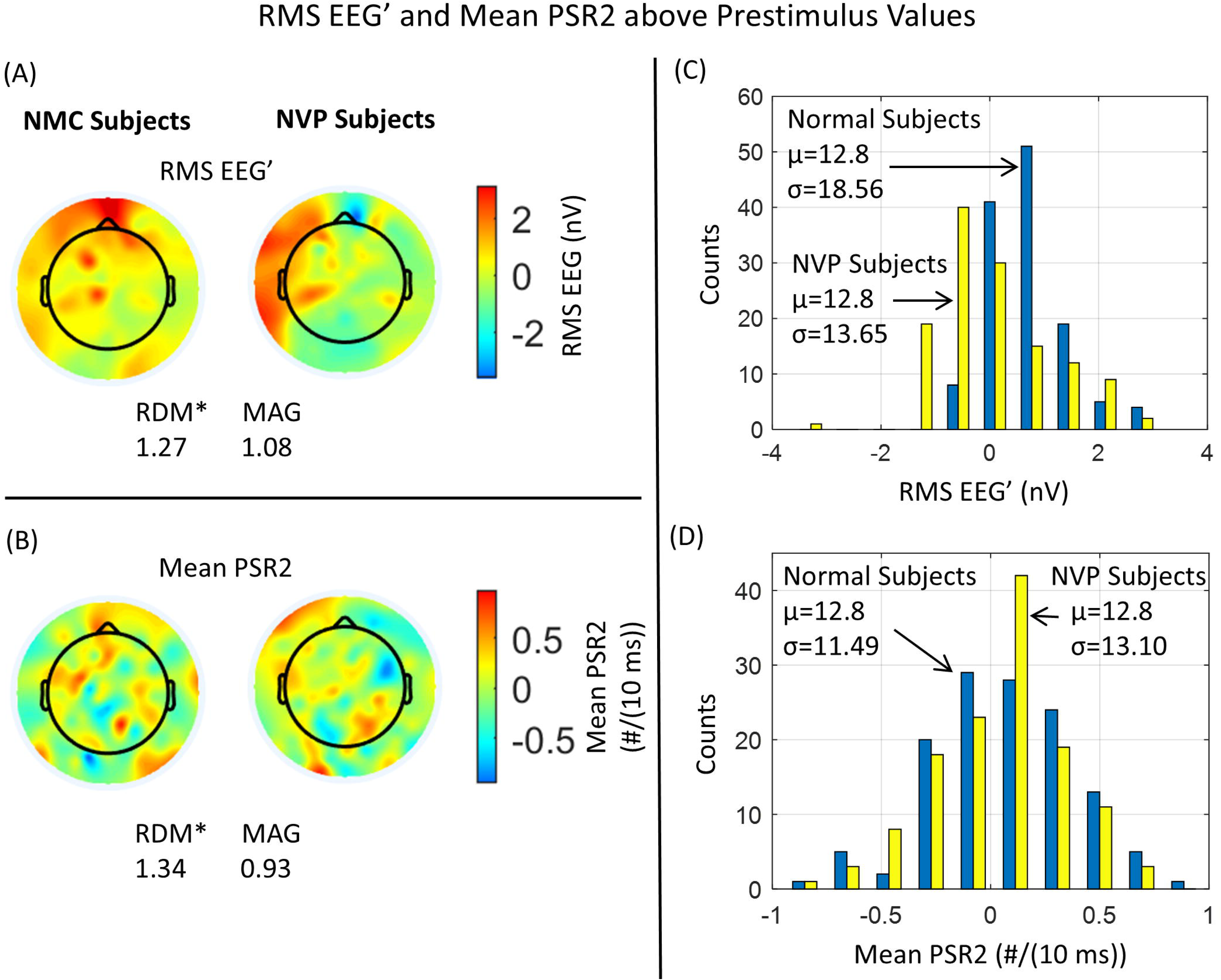
Activity patterns of the first derivative of EEG and PSR2 above baseline. (A) Spatial plots of RMS EEG’ above the baseline of NMC and NVP subjects, (B) corresponding spatial plots of mean PSR2 above the baseline, (C) histogram distribution of the RMS EEG’ values shown in spatial plots of A, and (D) histogram distribution of PSR2 values shown in spatial plots of B.

The spatial plots of PSR2 for the NMC and NVP subjects are given in Fig. 10B. For the NMC subject, the hot spots in PSR2 match fairly well with the hot spots in RMS EEG’. However, there are additional hot spots for the NVP subjects in the right parietal areas, which are absent in the RMS EEG’. A paired Student’s t-test failed to show any significant differences between the PSR2 values for the NMC and NVP subjects. However, the *RDM\** of 1.34 suggests that spatial distributions of PSR2 are significantly different from each other, as shown in Fig. 10B.

A histogram distribution of PSR2 is given in Fig. 10C. The mean values of histogram counts are the same for both groups, but the standard deviations, ó, are different. The ó is ±11.49 and ±13.10 for the NMC and NVP subjects, respectively. This translates to about a 14% difference between ó for the NMC and NVP subjects. The RMS values of histogram bin counts for NMC and NVP subjects are 16.81 counts and 17.84 counts, respectively. The RMS difference of histogram bin counts, RMS *H_diff_* is 5.58 counts. This translates to approximately a 16% difference in total bin counts, which could be considered a slight difference between the NMC and NVP subjects.

### D.4 Insight Moments of Object Recognition

EEG potentials in the broad band (3-49 Hz) and the corresponding phase slip rates in the alpha (7-12 Hz) at one of the electrodes in the right visual cortex are presented in Fig. 11. A choice was made to dispaly EEG potentials in the broad band because the P1, N1 and P2 peaks are easily recognizable which are difficult to see in the EEF filtered in the alpha (7-12 Hz) band. For reference, see Fig. 4 for NMC subjects and Fig. 6 for NVP subjects. The visual evoked potential peaks, i.e., P1, N1, and P2, during the stimulus period (0-240 ms) are related to the perception cycle when the subject looks at an object on the computer screen. The latency of these peaks, i.e., the delay between the start of the stimulus (0.0 s) and brain response, is very similar for the NMC and NVP subjects. For NMC subjects, these peaks are located at 122 ms, 190 ms, and 305 ms, respectively. For NVP subjects, these peaks are located at 115 ms, 214 ms, and 303 ms, respectively. Remarkably, these peaks for the two groups of subjects are very close to each other. Here, one group consists of eight NMC subjects, and the other consists of eight NVP subjects.

**Fig. 11.**
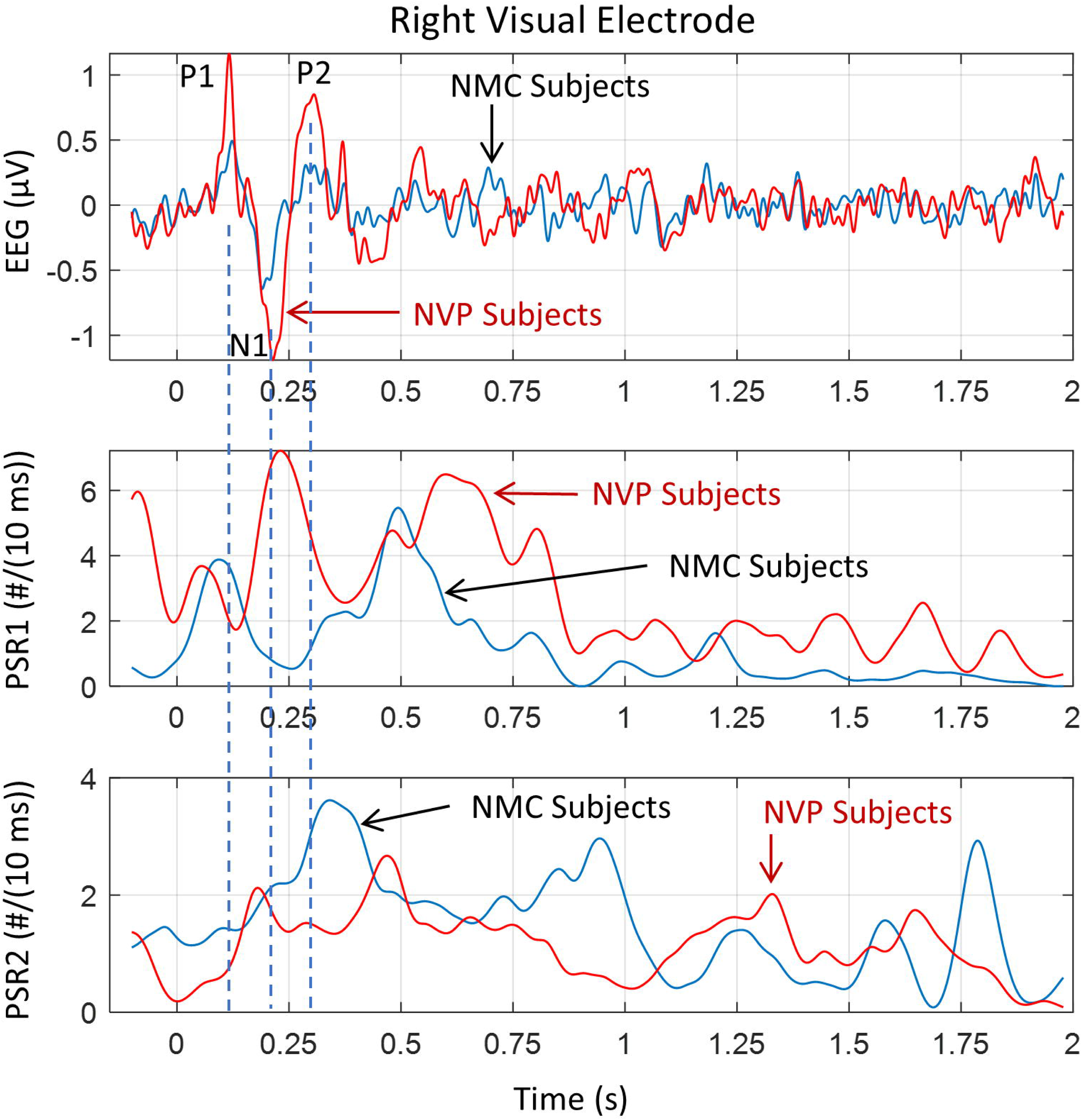
Visual evoked potentials and phase slip rates. EEG potentials in the broad (3-49 Hz) band and corresponding phase slip rates in the alpha (7-12 Hz) at one of the electrodes in the right visual cortex. (Top) EEG potentials with prominent P1, N1, and P2 peaks marked, (middle) corresponding PSR1, and (bottom) corresponding PSR2. Notice the subtle differences in EEG peaks and remarkable differences in PSR1 and PSR2 between NMC and NVP subjects.

During the poststimulus period (0.24 - 2.0 s), there are also prominent peaks in the EEG related to brain responses while the subject is analyzing and recognizing the name and form of the object [1]. These perception and decision-making processes extend from 0.0 s to about 1.2 s during the stimulus and poststimulus periods. After that, between 1.2 s and 2.0 s, brain activity begins to return to the regular background activity. Here, the background activity is related to a checkerboard pattern, which is displayed during the prestimulus and poststimulus periods. There are several peaks in the EEG during the poststimulus period of 0.0 to 1.2 s, which are at slightly different locations for NMC and NVP subjects, signifying that brain processes are slightly different for the two groups of subjects during the object recognition process.

The differences between the two groups of subjects are more pronounced when one looks at the temporal patterns of phase slip rates, viz., PSR1 and PSR2, given in the middle and the bottom plots of Fig. 11. The PSR1 is higher for NVP subjects as compared with NMC subjects, while PSR2 has the opposite trend. The PSR2 represents the net rate of number of neurons taking part in the genesis of EEG which one measures at the scalp or subdural electrodes. This suggests that the brain processes of the NMC subjects are changing more rapidly than those of the NVP subjects. One could probably say that the brain processes are more stable and gradually changing for NVP subjects as compared with NMC subjects.

As stated earlier, the visual evoked potentials, P1, N1, and P2, play a prominent role in the brain’s initial response to the object on the screen. Therefore, their spatial analysis is essential and is given in Fig. 12. There are noticeable differences in the spatial profiles of EEG for NMC and NVP subjects. The EEG potentials of P1 for NMC subjects show strong activity in a large area in the lower half of the plot, primarily in the visual, parietal, central, and left temporal regions. In comparison, for NVP subjects, it is mostly concentrated in visual, parietal, and central areas with a slight tilt toward the right side. The N1 positive potential activity for NMC subjects is distributed mainly in the left front and central areas, while for the NVP subjects, it is the left and right top portions of the plots. The negative components for both groups of subjects are distributed in the lower central and right areas. For NMC subjects, it is more focused, while for NVP subjects, it is distributed in larger areas in the lower right area of the plots. For the P2 potentials, both groups show some activity near the nose in the plots, which refers to the brain activity measured at electrodes on the forehead related to cortical activity in the frontal and orbital brain areas. The stark differences are evident in the lower posterior areas, where the NMC subjects exhibit less activity, while the NVP subjects show strong and distributed activity. There is a bright spot in the left frontal area for both groups of subjects.

**Fig. 12.**
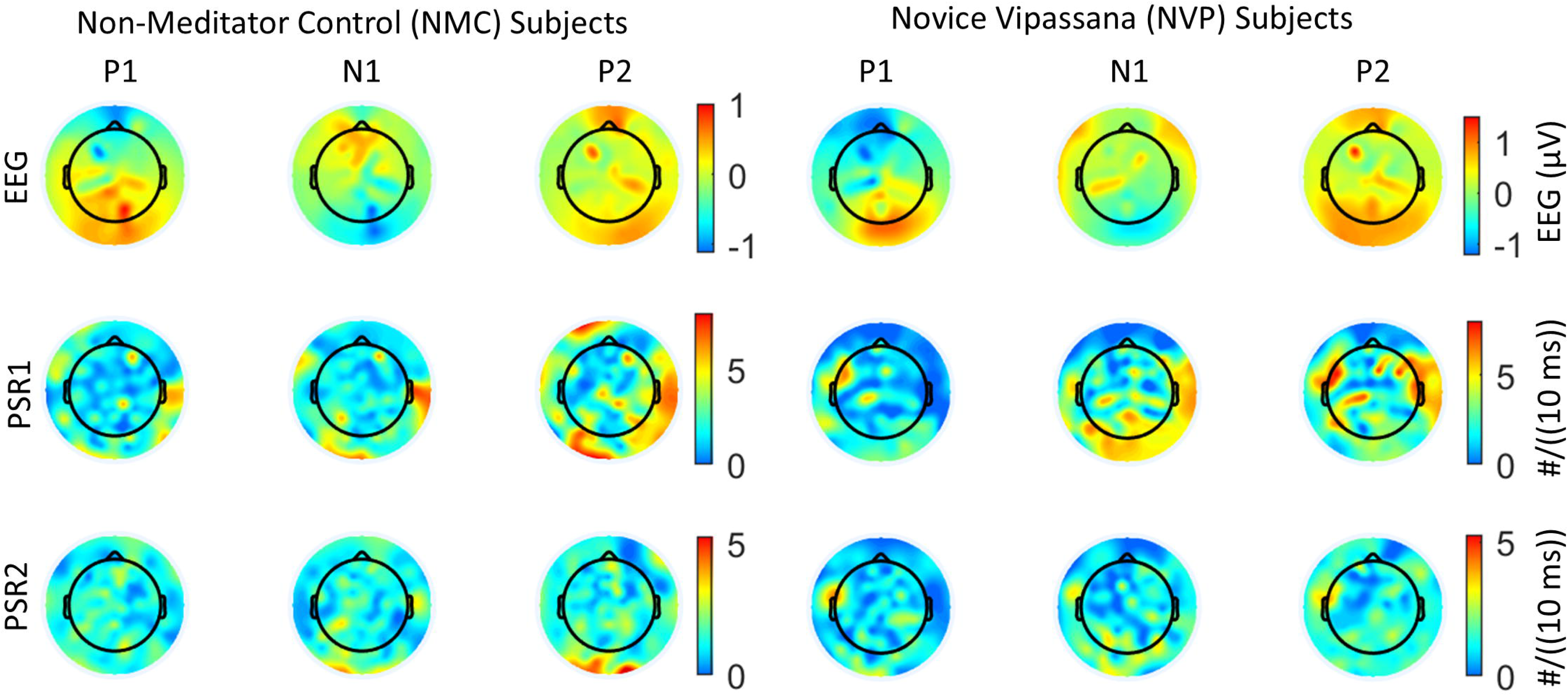
Spatial profiles of P1, N1, and P2 peaks. Spatial profiles of EEG, PSR1, and PSR2 for visual evoked response potentials, P1, N1, and P2 for NMC and NVP subjects. (Top row) EEG potentials in the broad (3-49 Hz) band, (middle row) PSR1, and (bottom row) PSR2. The PSR1 and PSR2 are in the alpha (7-12 Hz) band.

The spatial patterns for PSR1 (Fig. 12, middle row) are remarkably different from those of EEG potentials for all three peaks. For the P1 peak of NMC subjects, there is a bright spot near the right ear related to right temporal and auditory cortical brain activity. In contrast, for NVP subjects, there is a bright spot above the left ear, referring to the electrical activity of the left temporal and auditory cortices. Another noticeable difference in PSR1 plots is that NMC subjects have bright spots in the right frontal and central areas, which are missing for NVP subjects. This would suggest that there are some differences in the cortical activity of P1 peaks between NMC and NVP subjects. The PSR1 activity of the N1 peak is also different between NMC and NVP subjects. For the NMC subjects, it has bright spots near the right ear, right frontal, and left posterior central areas. In contrast, for the NVP subjects, PSR1 activity is distributed in the posterior region, near the right ear, and also exhibits some focused bright spots in the left central and parietal midline areas. There are also noticeable differences in the PSR1 plots of P2 peaks for NMC and NVP subjects. A noticeable difference is seen in the right front temporal and central posterior areas. The spatial plots of PSR2 have bright spots, which are well correlated with the bright spots in the PSR1 plots for all three peaks.

The *RDM\** and *MAG* values were computed for a comparative analysis between NMC and NVP subjects for all three peaks. These are listed in Table 2. Here, the reference model is the NMC subjects, and the test model is the NVP subjects. As stated earlier, if the reference and test models are the same, then *RDM\** will be zero and the *MAG* will be unity. From the values given in the table, it can be inferred that the EEG, PSR1, and PSR2 are significantly different for the NMC and NVP subjects across all three peaks.

**Table 2.** The *RDM\** and *MAG* values for NMC and NVP subjects at the P1, N1, and P2 peaks.

|  | P1 Peak |  | N1 Peak |  | P2 Peak |  |
| --- | --- | --- | --- | --- | --- | --- |
|  | *RDM\** | *MAG* | *RDM\** | *MAG* | *RDM\** | *MAG* |
| EEG | 0.486 | 1.585 | 0.976 | 0.866 | 0.673 | 2.290 |
| PSR1 | 0.849 | 0.943 | 0.656 | 1.430 | 0.687 | 1.025 |
| PSR2 | 0.768 | 0.880 | 0.705 | 0.985 | 0.635 | 0.869 |

### D.5 Time Frames of Insight Moments for NMC and NVP Subjects

There are several stages involved in recognizing the name and form of an object from memory [1], and our results indicate that there are subtle timing differences in these stages for NMC and NVP subjects. This is what we are investigating in here. These different stages of insight moments have been described in the literature [1, 36, 49, 43]. Stage I is the first impression of the object in the visual cortex, and it is called the “Awe” moment in the knowledge cycle. It is generally identified with the P1 wave of the visual evoked potentials. Stage II is related to the rapid exploration of the nature of the object in language and memory areas of the brain. The Stage III is the “Eureka” moment when one recognizes and identifies the object. The Stage IV is called the “Verification” stage [1] and it is related to the integration of the new knowledge in the memory for immediate future use, such as in visual object naming tasks. Stage V is the start of the return to the background activity of the brain. These different stages for a frontocentral electrode potentials are marked in Fig. 13A. The EEG potentials are in the 3-49 Hz band. Some of the prominent peaks are also marked, which are also shown in Fig. 3A and Fig. 11.

**Fig. 13.**
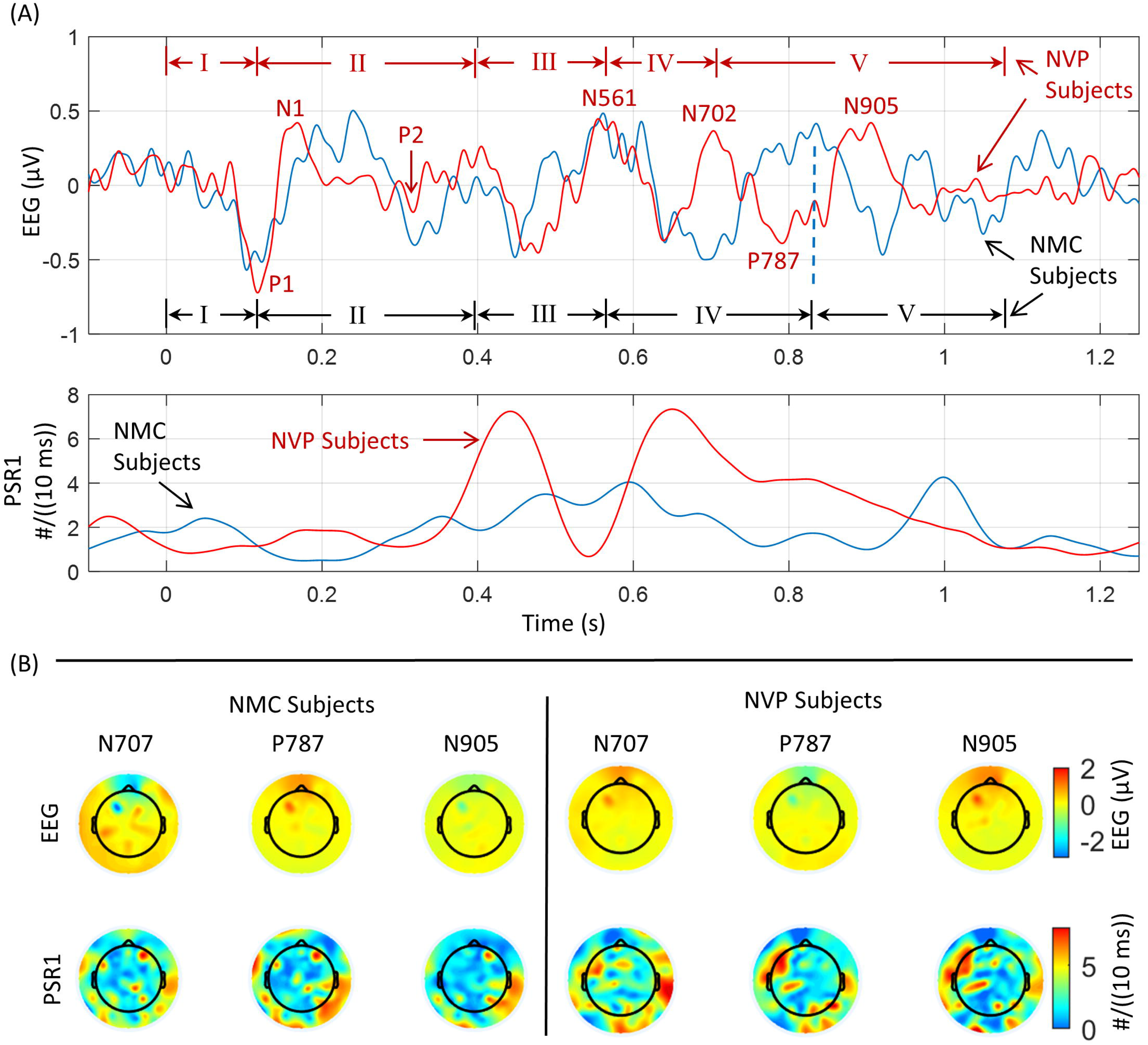
Insight moment stages in visual object recognition. (A) EEG potentials in the 3-49 Hz band for one of the electrodes in the frontocentral areas and the associated phase slip rates, PSR1. Different stages of the insight moments are also marked for NMC and NVP subjects. (B) Spatial plots of EEG and PSR1 of the prominent peaks for NMC and NVP subjects, and note their differences.

The initial first three stages of insight moments are similar, with only minor differences of a few milliseconds for NMC and NVP subjects. However, the duration of the fourth (IV) and fifth (V) stages is different. Stage I is from 0.0 s to the location of the P1 visual evoked potential. This will be the first impression of the object in the visual cortex. The endpoint of stage II is where the P2 peak ends, which can be well recognized in the visual evoked potentials given in Figs. 2 and 10. Stage III (305–561 ms) is the “Eureka” moment, when one recognizes and identifies the object [1, 36].

Stage IV is related to integrating the new knowledge into memory [1]. The Stage IV for the NVP subjects is from 561 to 702 ms, i.e., with a duration of 141 ms. In contrast, the Stage IV duration for NMC subjects is longer, ranging from 561 ms to 831 ms, with a total duration of 270 ms.

Stages III and IV are critical stages in the process of identifying and assimilating new knowledge into memory. NVP subjects seem to be quicker at assimilating new knowledge into memory than NMC subjects. This is also reflected in the bimodal shape of the PSR1 activity for NVP subjects (lower plot in Fig. 13A). *So, for NVP subjects, it takes 702 ms from the start of the stimulus to identify and assimilate the new knowledge into memory. In comparison, for the same task, it took 831 ms for NMC subjects.* Stage V marks the beginning of a return to background brain activity [1, 43].

The additional prominent peaks during the knowledge cycle, identified in Fig. 13A, are N707, P787, and N905. The spatial plots of EEG (3-49 Hz) potentials and their respective PSR1 are given in Fig. 13 B. The spatial plots of the EEG potentials for NMC and NVP subjects are significantly different (p < 0.01), which was confirmed by the paired Student’s t-test. Spatial differences are also noticeable for these three peaks between NMC and NVP subjects. For the N707 peak, the RDM* and MAG between NMC and NVP subjects were: 0.697 and 1.19. The same comparative values for the peak P787 were: 0.787 and 1.03. For the peak N905, these were: 0.85 and 1.29. These values suggest that the spatial plots of PSR1 for the NMC and NVP subjects are significantly different from each other.

## E. Discussions

Our primary objective was to quantify the differences in decision-making processes between NMC and NVP subjects. Our results demonstrate quantifiable differences in four biomarkers that we utilized in this study. The temporal and spatial patterns of EEG, EEG’, PSR1, and PSR2 indicate that different brain regions are activated in NMC and NVP subjects. Another remarkable finding is that NVP subjects are quicker (702 ms) compared to NMC subjects (831 ms) in completing the visual task of identifying and assimilating new knowledge into memory. This is a remarkable finding that suggests people can be trained to perform tasks more efficiently for visual object recognition. The emphasis here is on visual object recognition processes, which are commonly employed in various meditation techniques. This finding not only suggests potential applications in other meditative techniques but also underscores the need for further research in this field. It also suggests that similar results may be observed in other tasks, such as motor function-related tasks, which should be examined through carefully designed studies comparing non-meditator control subjects with those who have undergone meditative training in those particular tasks. An example of motor function-related tasks is the use of hand gestures, i.e., Mudras, which are very common in meditation practices.

Although many studies have been conducted to examine the efficacy of Vipassana in the context of mindfulness-based interventions, relatively few studies have focused on traditional Vipassana practitioners. Prior work from our group has examined EEG profiles of three proficiency-linked groups of Vipassana meditation practitioners while they engaged in a structured sequence of meditation practice (Anapana or focused attention, Vipassana or mindfulness, and Metta or loving-kindness) and found that long-term meditators (seniors and teachers) showed widespread trait and state differences as compared to novice meditators [28]. Examination of P300 dynamics in the same three groups of meditators during the performance of the ANGEL task had revealed significant state-trait influences of meditation practice [29]. Future work could therefore evaluate the applicability of the four biomarkers employed in the current study across distinct meditation states in NVPs and in long-term meditators.

Our work with novice Vipassana-trained subjects was facilitated by the availability of their data sets and can be readily applied to other datasets from Vipassana meditators [50]. However, the techniques and tools developed for this work are generic and versatile, and can be potentially applied to any EEG data set to augment the existing set of measures. For instance, these tools can be used to study the behavior of default mode networks in the spontaneous EEG or evoked potential studies [51]. Similarly, our tools are applicable across diverse meditation practices, including Zen, Mindfulness, Insight, other Buddhist and non-Buddhist traditions [52, 53, 54, 55]. Indeed, recent work examining EEG data from four different meditation traditions revealed both a common core and tradition-specificity in the neurodynamic profiles [56]. When the mind is trained to remain stable and focused, tools from our current work can augment traditional measures in detecting subtle changes in brain activity from EEG data [57, 58]. Their sensitivity extends to calm-abiding practices that cultivate stability and insight (Samatha & Vipassana) and to practices that foster compassion, generosity, morality, joyous effort, and loving-kindness [59, 60, 61]. We hope that other investigators will adopt these tools in their research work.

We have used four biomarkers in this study. The spatial profiles of EEG and EEG’ are similar, with some minor differences which are not significant. Similarly, the spatial differences in the PSR1 and PSR2 plots are also minor. However, there are some significant differences between the spatial plots of EEG and its related PSR1, as well as between EEG’ and PSR2, that require explanation. Refer to Figs. 6 and 8. We first introduced these biomarkers in our recent work [4], where we mentioned that the first-order derivative of the EEG data removes linear trends and amplifies nonlinear trends and oscillatory activity in the data. The computation of the phase from EEG or EEG’ by use of the Hilbert transform does not depend upon the amplitude of the data but rather on large and small perturbations in the data. Even at the small perturbations in the data, a ±ð phase shift is introduced, which could influence the computations of PSR1 and PSR2 at numerous spatial locations. Additionally, within a given time slice of the EEG (or EEG’) data, large amplitude values may occur on a few channels, while small perturbations in the data may occur on other channels. Thus, the spatial locations of PSR1 and PSR2 could be different from the spatial locations of EEG or EEG’, respectively. These could be the reasons why the EEG and PSR1, as well as EEG’ and PSR2, show some significant differences in the spatial plots. One could verify these by source location analysis from EEG and PSR1, or from EEG’ and PSR2. This is a significantly big task, perhaps suitable for a separate study.

Our results, in line with earlier research [4], underscore the robustness of phase slip activity across broader brain areas, even when EEG potentials are low or near the zero-crossing lines. This robustness suggests that phase slip rates could be a more sensitive biomarker than EEG or EEG’ for detecting changes in brain activity. Importantly, this could significantly enhance our understanding of brain functions, particularly in scenarios such as the study of the EEG of anesthetized patients [62], the EEG data of subjects engaged in the Tukdum practice of Buddhist meditation near the time of death [63], or the EEG of subjects going through near-death experiences [64]. In these situations, the use of phase slip rate biomarkers derived from EEG, as well as the first and second-order derivatives of EEG, could provide valuable insights into brain activity, where EEG alone may not be sensitive enough to detect widespread changes in brain activity [4]. Furthermore, phase slip rates could serve as a sensitive gauge to monitor the level of anesthesia or the patient’s level of consciousness during surgery or meditation.

Here, we have limited our work to the alpha (7-12 Hz) band only, which, in general, is the most predominant band in the power spectrum and also plays a crucial role in visual object recognition and insight moment analysis studies [1]. In future studies, this work could be extended to other EEG bands, such as theta, alpha, beta, and gamma bands, which may provide additional insights into the similarities and differences between NMC and NVP subjects in decision making processes. Additionally, phase amplitude couplings between different bands, such as theta-alpha or beta-gamma bands, may be another area to explore.

In summary, we demonstrated that cortical phase transitions alter during decision making and that brain functions differ between non-meditators and novice Vipassana meditators. These methods and results may aid future experimental designs in visual cognition and mindfulness research, as well as contribute to educational neuroscience and the development of neurofeedback technologies for clinical applications. This is our hope.

## F. Data availability statement

Publicly available data sets were used in this study. This data can be found at: https://osf.io.

## G. Author contributions

RJK: Conceptualization, Methodology, Writing – original draft, Writing – review & editing. AKN: Data curation, Data analysis, Writing – original draft, Writing – review & editing.

AS: Data curation, Data analysis, Writing – review & editing. RV: Data Curation.

PNR: Project administration, Supervision, Writing – review & editing. BK: Project administration, Supervision, Writing – review & editing.

CR: Conceptualization, Methodology, Data curation, Data analysis, Software, Visualization, Writing – original draft, Writing – review & editing.

## Acknowledgment

Professor Jens Hauesien from the Technical University of Ilmenau, Germany, made several technical and grammatical suggestions to improve the manuscript’s writing. We want to thank him for his efforts.

## Abbreviations

EEG: Electroencephalogram or Electroencephalography
EEG’: First order derivative (*d/dt*) of EEG
NMC: Non-Meditator Controls
NVP: Novice Vipassana
RDM*: Relative Difference Measure
PSR: Phase Slip Rate or Phase Slip Rates
PSR1: Phase Slip Rate extracted from EEG
PSR2: Phase Slip Rate extracted from EEG’
Poststimulus+: Combined duration of stimulus and poststimulus period

